# Palaeoproteomic insights into the deep roots of the cave bear lineage in Europe

**DOI:** 10.1101/2025.10.05.680322

**Authors:** Amanda Gutiérrez-Carbajal, Elena Santos, Esther Lizano, Tomas Marques-Bonet, Ricardo Fong-Zazueta, Aurora Grandal-d’Anglade, David M. Alba, Ana García-Vázquez, Asier Gómez-Olivencia, Mónica Villalba de Alvarado, Luca Pandolfi, Lorenzo Rook, Montserrat Sanz, Juan Luis Arsuaga, José María Bermúdez de Castro, María Martinón-Torres

**Affiliations:** Departament de Medicina i Ciències de la Vida, Institut de Biologia Evolutiva (CSIC-UPF), Universitat Pompeu Fabra, Barcelona, Spain; Centro Mixto UCM–ISCIII de Evolución y Comportamiento Humanos Madrid, Spain; CENIEH (Centro Nacional de Investigación sobre la Evolución Humana), Burgos, Spain; Cátedra de Otoacústica Evolutiva y Paleoantropología (HM Hospitales-Universidad de Alcalá), Departamento de Ciencias de la Vida, Universidad de Alcalá, Alcalá de Henares, Madrid, Spain; Institut Català de Paleontologia Miquel Crusafont (ICP-CERCA), Universitat Autònoma de Barcelona, Cerdanyola del Vallès, Barcelona, Spain; Unidad de Paleobiología, ICP-CERCA, Unidad Asociada al CSIC por el IBE UPF-CSIC, Cerdanyola del Vallès, Barcelona, Spain; Catalan Institution of Research and Advanced Studies (ICREA), Passeig de Lluís Companys, Barcelona, Spain; CNAG, Centro Nacional de Analisis Genomico, Barcelona, Spain; Département de sciences biologiques, Université de Montréal, Montréal, Canada; Instituto Universitario de Xeoloxía, Universidade da Coruña, ESCI, A Coruña, Spain; ArchaeoScience Platform (ASp), Research Institute of the University of Bucharest (ICUB), University of Bucharest, Bucharest, Romania; Departamento de Geología, Facultad de Ciencia y Tecnología, Universidad del País Vasco–Euskal Herriko Unibertsitatea (UPV/EHU), Leioa, Spain; Sociedad de Ciencias Aranzadi, Donostia–San Sebastián, Spain; Departamento de Prehistoria, Historia Antigua y Arqueología, Universidad Complutense de Madrid, Madrid, Spain; Musée de l’Homme, Paris, France; Dipartimento di Scienze della Terra, Università di Pisa, Pisa, Italy; Dipartimento di Scienze della Terra, Paleo[Fab]Lab, Università degli Studi di Firenze Firenze, Italy; Grup de Recerca del Quaternari (GRQ)-SERP, Departament d’Història i Arqueologia, Universitat de Barcelona, Barcelona, Spain; Faculdade de Letras, UNIARQ-Centro de Arqueologia da Universidade de Lisboa, Lisboa, Portugal; Departamento de Geodinámica, Estratigrafía y Paleontología, Facultad de Ciencias Geológicas, Universidad Complutense de Madrid, Madrid, Spain; Department of Anthropology, University College London, London, UK

**Author notes:** Corresponding author (A.G.-C.). **Author Contributions:** Conceptualization: M.M.-T. and E.S.; Data curation: A.G.-C.; Formal analysis: A.G.-C., Methodology and analysis: A.G.-C., with assistance and guidance from M.M.-T., E.S., T.M.-B., and E.L., and with specific contributions from R.F.-Z.; Paleontological material contribution: E.S., D.M.A., A.G.-O., A.G.-A., L.R., L.P., M.S., A.G.-V., M.V.A. and J.L.A.; Writing original draft: A.G.-C.; Writing-review and editing: A.G.-C. with input from all coauthors; Supervision: M.M.-T., E.S., and T.M.-B.; Project administration and funding acquisition: J.L.A., M.M.-T. **Competing Interest Statement:** The authors declare no competing interests.

**Keywords:** Palaeoproteomics, Cave Bears, Pleistocene, Enamel, Proteins

## Abstract

Palaeoproteomics has emerged as a powerful tool for reconstructing the evolutionary history of extinct species, particularly when ancient DNA is poorly preserved or beyond recovery. Here, we present the first large-scale enamel proteomic study focused on the cave bear, spanning specimens from the Early to Late Pleistocene. A primary objective was to determine whether the ursid population from level TD4 of Gran Dolina (Sierra de Atapuerca, Spain) belongs to the cave or the brown bear lineage, a long-standing taxonomic debate. We analyzed specimens from the Atapuerca sites, alongside comparative material from other southwestern European localities. Using LC-MS/MS and an acid demineralization protocol without enzymatic digestion, we successfully recovered enamel proteomes from all fossil samples, including the oldest specimens. Protein profiles were obtained for each extinct ursid, enabling the identification of taxonomically informative peptides across multiple individuals per taxon. Notably, two novel single amino acid polymorphisms (SAPs), found in ameloblastin (AMBN) and alpha-1 antitrypsin (SERPINA1), were restricted to Middle and Late Pleistocene cave bears and may represent new phylogenetic markers for this clade. This study provides the first molecular phylogenetic placement of *Ursus dolinensis*, supporting its basal position within the speloid lineage, consistent with its proposed ancestral status. Our results highlight the strong phylogenetic signal preserved in dental enamel and the exceptional biomolecular preservation at Atapuerca, providing a robust framework for reconstructing the evolutionary history of Ursidae. Moreover, the consistent recovery of systemic proteins, such as serpins, underscores the potential of enamel proteomes to capture not only evolutionary relationships but also physiological signals relevant to extinct ursids.

**Significance:** The phylogenetic position of *Ursus dolinensis* within the Ursidae family has been a subject of long-standing debate. While some authors place it within the speloid lineage, others suggest affinities with arctoid bears. Here, we analyze the bear population from level TD4 of Gran Dolina (Sierra de Atapuerca, Spain) using palaeoproteomic methods for the first time. Molecular evidence supports the inclusion of *U. dolinensis* within the speloid lineage, in a basal position, supporting its potential ancestral position. These findings extend the evolutionary depth of the speloid clade in Europe and demonstrate the power of enamel palaeoproteomics to resolve deep-time relationships beyond the limits of ancient DNA, revealing the deep evolutionary roots of the lineage.

## Introduction

The evolutionary history of cave bears (the speloid lineage) has long been a subject of extensive paleontological and genetic interest due to their extensive fossil record and ecological significance during the Pleistocene. These ursids, distributed from the Iberian Peninsula across Eurasia to northeastern Siberia [1], are predominantly known from karstic deposits, reflecting their hibernation behavior [2–4]. Phylogenetically, cave bears (*Ursus deningeri*-*Ursus spelaeus* sensu lato) form a sister clade to the extant brown (*Ursus arctos* Linnaeus, 1758) and polar bears (*Ursus maritimus* Phipps, 1774), diverging approximately 1.2–1.6 million years ago (Ma) based on nuclear and mitochondrial DNA clocks [5, 6]. The lineage persisted until ∼24–26 thousand years ago (ka), when the last representatives became extinct, likely following a demographic decline possibly influenced by climatic changes and human competition for cave habitats [7, 8].

Multiple hypotheses have been proposed regarding the origin and diversification of this lineage. Morphological studies, ranging from broad biochronological syntheses to population-level and morphometric analyses, suggest either gradual (anagenetic) evolution or more complex branching scenarios [9–12]. *Ursus etruscus* Cuvier, 1823 is often regarded as the common ancestor of both cave and brown bears [2, 3, 13]. Ancient DNA (aDNA) analyses have clarified relationships among Late Pleistocene cave bear taxa—*Ursus spelaeus* Rosenmüller, 1794, *Ursus kanivetz* Vereshchagin, 1973, and *U. kudarensis* Baryshnikov, 1985—revealing regional structuring and divergence [14]. However, preservation limits usually restrict genomic data to the Middle Pleistocene or younger [15, 16]. The oldest mitochondrial sequences derive from *Ursus deningeri* von Reichenau, 1904 specimens at Sima de los Huesos (∼430 ka) [17, 18], consistent with recent cranial evidence that situates the hominins within this chronological range [19], leaving earlier phases of speloid evolution poorly resolved. This gap is particularly evident for *Ursus dolinensis* García & Arsuaga, 2001 from the TD4 level in the Gran Dolina site (Sierra de Atapuerca, Spain). While some authors position *U. dolinensis* at the base of the speloid lineage [20], others argue for affinities with archaic arctoid forms, citing its mosaic of primitive and derived traits [13, 21, 22]. Therefore, alternative biomolecular approaches are necessary to probe the evolutionary history of the cave bear lineage more deeply.

Palaeoproteomics offers a means to bridge this gap. Proteins, being more chemically stable than DNA, can persist for over a million years in mineralized tissues such as dental enamel [23, 24]. Advances in mass spectrometry and bioinformatics now allow recovery of phylogenetically informative single amino acid polymorphisms (SAPs) from fossil proteomes [25]. This approach has already illuminated aspects of Pleistocene mammals [26] and hominin evolution [27-29], yet carnivores—and ursids in particular—remain largely unexplored. Here, we present the first comprehensive enamel proteomic analysis of fossil bears across a broad temporal and geographic range, spanning the Early to Late Pleistocene. By analyzing proteomes from *U. dolinensis, U. deningeri*, and *U. spelaeus* and integrating them with extant *Ursus* and *Ailuropoda* references, we aim to (1) characterize enamel proteome composition and (2) evaluate its phylogenetic utility for clarifying cave bear evolutionary history.

## Results

### Proteomic Recovery

Ancient peptides were successfully recovered from all the analyzed 55 fossils, with peptide-spectrum matches (PSMs) varying across temporal groups. Recovery was generally lower in Early Pleistocene and open-air specimens, whereas Middle Pleistocene samples—particularly those from Sima de los Huesos—reached counts comparable to Late Pleistocene individuals (Figure 1.A). An extant specimen of *U. arctos* used as a control yielded fewer PSMs overall, although relative intensities of the peptides remained within the expected range (***SI Supplementary Results, Fig. S4***). Across fossils, fragment length distributions were dominated by short peptides (8–10 amino acids), consistent with long-term hydrolysis. Middle Pleistocene specimens produced the highest peptide counts (up to 390), while Late Pleistocene individuals preserved a larger proportion of longer sequences (>15 amino acids) (Figure 1.B). Early Pleistocene specimens exhibited more advanced degradation, with lower yields and shorter fragments. Post-translational modifications, particularly extensive deamidation, confirmed the antiquity of all fossil sequences, with values consistently higher than in the modern control (Figure 1.C). Oxidation patterns showed no marked differences.

**Figure 1.**
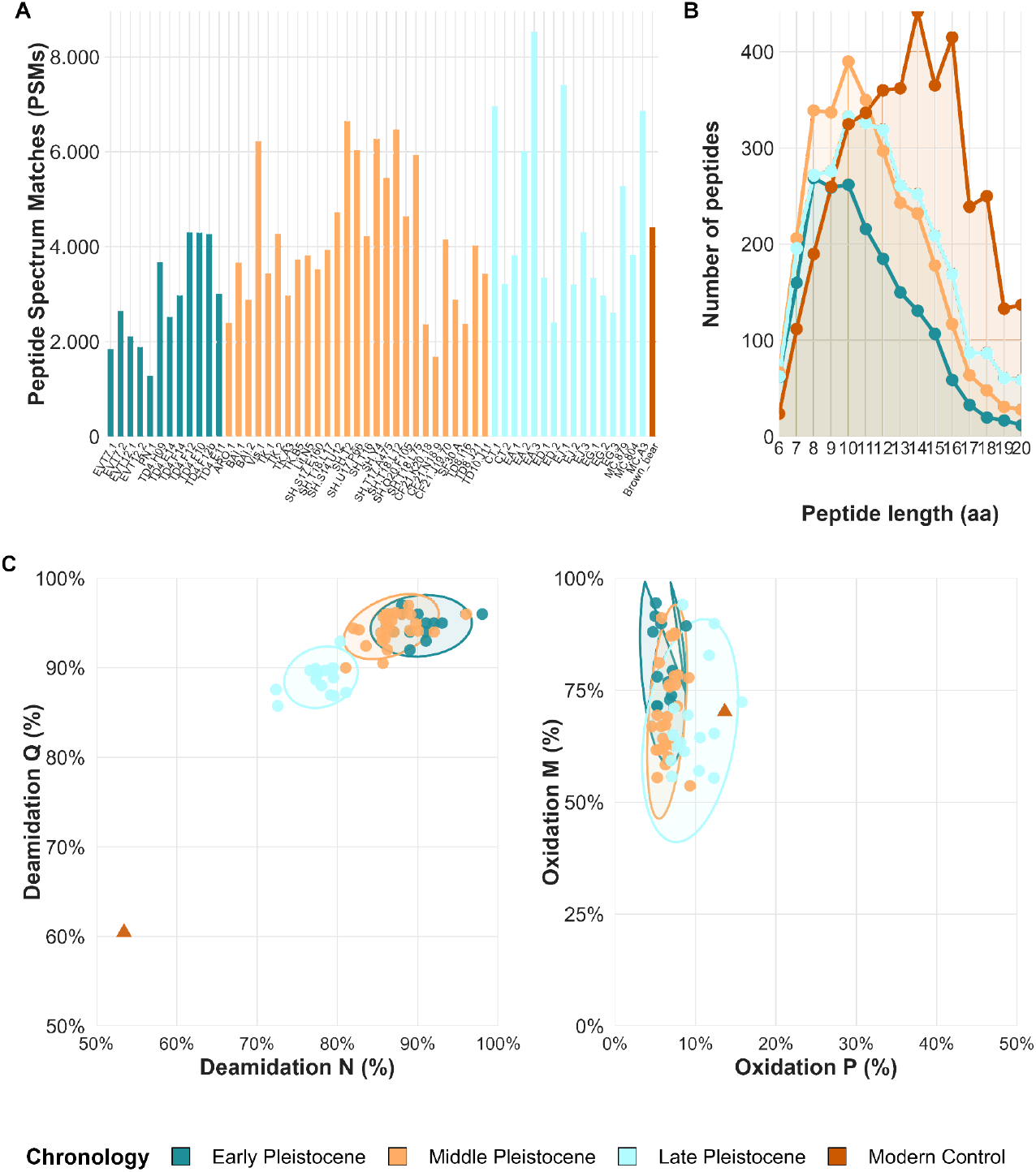
Endogenous origin and preservation of enamel peptides across temporal groups. **A**. Proteomic recovery measured as peptide-spectrum matches (PSMs) per specimen. **B**. Distribution of peptide lengths. **C**. Mean rates of deamidation at asparagine/glutamine (N/Q) and oxidation at methionine/proline (M/P).

Between 10 and 16 proteins were confidently identified per specimen. Enamel-specific proteins—including amelogenin (AMELX/AMELY), enamelin (ENAM), ameloblastin (AMBN), amelotin (AMTN), odontogenic ameloblast-associated protein (ODAM), and matrix metalloproteinase-20 (MMP20)—were consistently recovered, often accompanied by plasma proteins such as albumin (ALB), alpha-2-HS-glycoprotein (AHSG), alpha-1-antitrypsin (SERPINA1), antithrombin III (SERPINC1), and occasionally by collagen types I and XVII (COL1A1, COL1A2, COL17A1). Less frequent identifications included apolipoprotein A1 (APOA1) and zinc finger protein (ZFC3H1). The combined use of two search engines improved detection but yielded largely overlapping results.

Amelogenin isoforms enabled sex determination in most individuals: 27 were identified as male (AMELY-specific peptides), 25 as female or likely female (AMELX-only), and 3 remained inconclusive. Complete recovery metrics and platform-specific comparisons are provided in ***SI Dataset S2– Peptide Recovery and Identification***.

### Sequence Variation

Protein alignments reveal high overall sequence conservation across ursids, yet eight proteins—AHSG, AMBN, ALB, COL17A1, ENAM, MMP20, SERPINA1, and ZFC3H1—contained phylogenetically informative single amino acid polymorphisms (SAPs). In total, 48 SAPs were identified. AMBN and ALB had the highest counts (11 each), followed by COL17A1 and ENAM (9 each). Most variants distinguished *Ursus* from *Ailuropoda* (63%), while 27% represented intra-*Ursus* variation, and approximately 10% appeared potentially unique to extinct cave bear lineages (***SI Supplementary Results Table S1-Figure S4***). Only three high-confidence SAPs passing stringent validation criteria were retained.

Two novel variants, not previously documented in the reference proteomes of ursids, were detected in AMBN and SERPINA1. A substitution of tyrosine (Y) to serine (S) at position AMBN-249 was consistently present in *U. deningeri* and *U. spelaeus* specimens from multiple sites in Spain, Portugal, and Italy (Figure 2). Two Middle Pleistocene individuals from the level TD8 (Gran Dolina, Spain) and Aroeira (Portugal) presented ambiguous signals, suggesting incomplete fixation of this variant or potential heterozygosity (details in ***SI Supplementary Results***). A second AMBN substitution, leucine (L) to valine (V) at AMBN-278, was observed in a single specimen from Sala Fantasma and is reported cautiously. Both AMBN variants occur in exon 13, a variable region among mammals, and likely represent conservative changes [30]. In SERPINA1, an asparagine (N) to threonine (T) substitution at position 341 was consistently recovered in *U. deningeri* and *U. spelaeus* as well as in one *U. arctos* individual from Monte Cucco (Italy), and absent in *U. dolinensis*, providing an additional marker for the speloid lineage.

**Figure 2.**
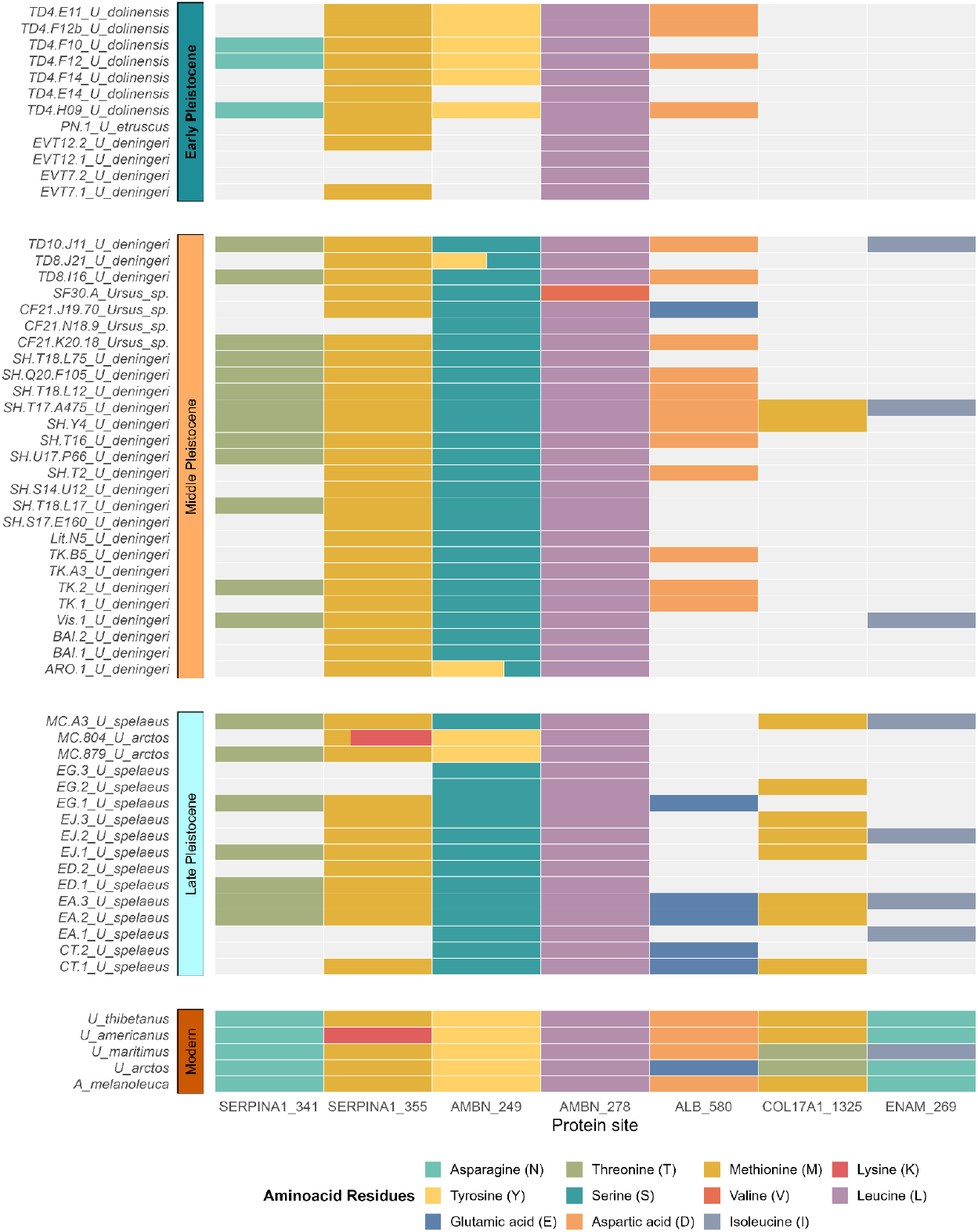
Sequence variation across enamel and serum proteins in fossil and modern bears. Color-coded matrix displaying informative amino acid positions. Columns represent variable sites and rows correspond to individual specimens. Partial or split-colored bars indicate heterozygous positions, where two different amino acids were detected in the same protein from a given specimen.

Other non-unique SAPs also proved informative. At ALB-580, *U. dolinensis* and *U. deningeri* shared the ancestral state with several extant ursids, while *U. spelaeus* and *U. arctos* carried the derived variant. At ENAM-269, isoleucine (I) occurred in *U. deningeri, U. spelaeus*, and *U. maritimus*, contrasting with asparagine (N) in other bears. In COL17A1-1325, *U. deningeri* and *U. spelaeus* shared methionine (M), whereas *U. arctos* and *U. maritimus* carried threonine (T). Coverage for these sites was incomplete in *U. dolinensis*, preventing a complete assessment. Collectively, these patterns reveal conserved molecular cohesion within the *U. deningeri–U. spelaeus* clade, retention of ancestral states in *U. dolinensis*, and several lineage-specific markers with potential for biochronological and phylogenetic applications.

### Phylogenetic reconstruction

Phylogenetic analyses of enamel proteomes yielded congruent topologies across Bayesian and maximum likelihood methods (Figure 3). All reconstructions recovered *Ursus* as monophyletic, with a basal divergence between arctoid bears and the speloid lineage. Within speloids, *U. dolinensis* was consistently recovered as the earliest diverging member, preceding *U. deningeri* and *U. spelaeus*. Although statistical support for this placement was modest, the topology remained stable across datasets. *Ursus deningeri* and *U. spelaeus* formed a strongly supported clade, confirming their close evolutionary relationship and their more derived status relative to *U. dolinensis*. Bayesian inference produced the best resolved tree, with consistent placement of both extinct and extant taxa.

**Figure 3.**
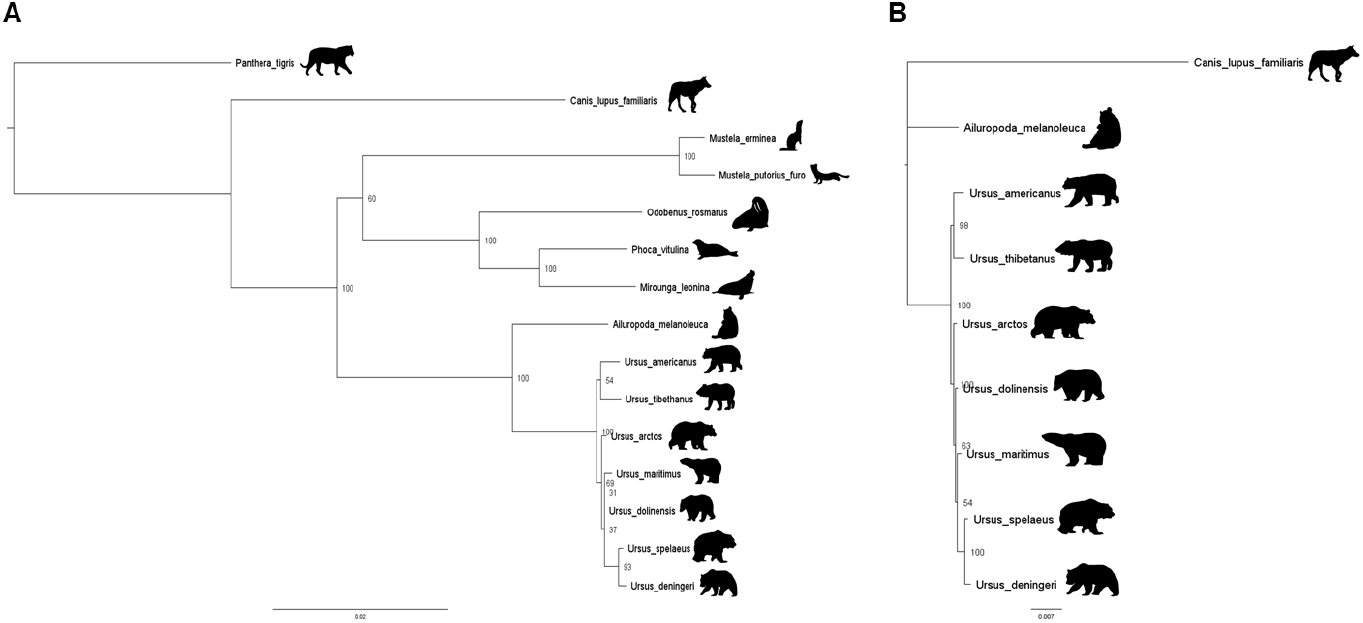
**A**. Maximum likelihood phylogenetic tree of selected Carnivora, including cave bears, inferred from enamel protein sequences using IQTree (Carnivora Dataset). **B**. Bayesian phylogenetic tree of the Ursidae based on concatenated enamel protein sequences (Ursidae dataset), inferred with MrBayes under a mixed model across 3,000,000 generations.

An unexpected result was the placement of *U. maritimus* as sister to the *U. deningeri–U. spelaeus* clade instead of clustering with *U. arctos*, as reported in previous studies. This position was weakly supported and may reflect either database limitations or convergent molecular signals. Additional analyses and full statistical details are provided in the ***SI Supplementary Results. Expanded Phylogenetic Trees and Node Support***.

## Discussion

### Enamel Proteome Composition

This study provides the first large-scale characterization of enamel proteins in the cave bear lineage. Across all 55 specimens, enamel-specific proteins—AMELX, AMBN, AMTN, and ENAM—were consistently identified, confirming their stability and central role in enamel formation, consistent with reports from other Pleistocene studies [26]. Plasma proteins, such as ALB and AHSG, were also commonly recovered; the latter appeared in 43 specimens, supporting its proposed endogenous incorporation into enamel [28]. By contrast, TUFT1 and KLK4, well-known components of mammalian enamel [31], were largely absent, suggesting either taxon-specific differences or selective degradation patterns. MMP20 was consistently recovered and exhibited sequence variation, reinforcing its relevance in enamel proteome studies. Collagen peptides were abundant in Late Pleistocene *U. spelaeus* but scarce in older specimens, and no informative variants were observed in COL1 genes, consistent with their strong evolutionary conservation [32]. ZooMS analyses have similarly reported no COL1 markers distinguishing *U. spelaeus* from *U. arctos* [33]. In contrast, COL17A1 was detected in nearly all specimens and showed moderate, non-lineage-specific variation. Its essential role in enamel formation, demonstrated by defects in COL17A1-deficient mice [34], underscores both its preservation potential and comparative relevance across taxa.

Particularly noteworthy is the recurrent recovery of three serine protease inhibitors (serpins)—alpha-1-antitrypsin (SERPINA1), antithrombin III (SERPINC1), and, more sporadically, pigment epithelium-derived factor (SERPINF1). Specifically, SERPINA1 was detected in nearly all specimens, often with high peptide counts and conserved post-translational modifications, including a lineage-specific SAP. Their consistent presence in enamel proteomes raises questions about the pathways by which they became incorporated. One possibility is a role in amelogenesis, given their co-occurrence with proteases such as MMP20 and KLK4 and previous evidence for interactions with mineralized tissues [35, 36]. Alternatively, they may reflect systemic physiology, particularly hibernation, which in bears involves metabolic suppression, cardiovascular regulation, and immune modulation [37–40]. Studies in extant hibernators support a role of circulating serpins in these processes [41–43], making their frequent recovery in cave bears particularly intriguing. If enamel mineralization overlapped with torpor, serpins transiently elevated in circulation could have been incorporated into the developing matrix.

While additional comparative data from other carnivorans are needed, the evidence suggests that enamel proteomes capture not only phylogenetic signals but also systemic proteins potentially linked to physiology. This dual dimension highlights the broader scope of palaeoproteomics for investigating both evolutionary and ecological aspects of extinct species. Expanding comparative analyses across both modern and fossil carnivorans will be critical to assess whether these serpins represent lineage-specific traits or more general physiological patterns. Although serpins have been sporadically reported in dental proteomes of non-ursid mammals [31, 44, 45], the frequent detection of SERPINA1 in cave bears points to a distinctive feature of the genus *Ursus*.

### Phylogenetic Relationships within the Cave Bear Lineage

Our palaeoproteomic analyses refine phylogenetic relationships within the cave bear lineage. The placement of *U. dolinensis* as basal relative to *U. deningeri*-*U. spelaeus* clade supports its interpretation as an early member of the speloid clade, shortly after ts divergence from arctoid bears, consistent with some morphology-based phylogenetic hypotheses [20]. Within speloids, *U. deningeri* and *U. spelaeus* share two derived substitutions (AMBN-249 and SERPINA1-341), confirming their genetic cohesion [17, 18]. Their minimal proteomic distinction, restricted to a single SAP (ALB-580), highlights both their close relationship and the limited sequence divergence detectable in enamel proteins. The possibility that *U. deningeri* represents a direct ancestor of *U. spelaeus* cannot be formally evaluated within the dichotomic phylogenetic framework applied here, although this scenario remains consistent with the molecular continuity observed.

Comparisons with extant taxa broadly align with genomic studies: *U. arctos* emerges as the closest living relative of cave bears [16, 46], while other ursids cluster as expected. The only exception is *U. maritimus*, which is occasionally grouped with the *U. deningeri–U. spelaeus* clade instead of *U. arctos*. This placement was weakly supported and likely reflects either database incompleteness or convergent signals, possibly linked to the shared SAP at ENAM-269 (I) observed in *U. maritimus* and several Middle to Late Pleistocene specimens (see ***SI Supplementary Results***). The lack of peptide coverage at this site in Early Pleistocene individuals, however, limits further evaluation.

Our results illustrate both the utility and the limits of enamel proteomes for phylogenetic inference. While major clades are consistently resolved, species-level distinctions remain challenging due to high sequence conservation, incomplete coverage, and the fragmentary nature of ancient proteins [47].

### Proteomic Signatures and Evolutionary Dynamics

Beyond clarifying phylogenetic placement, enamel proteomes reveal evolutionary signatures across both time and space. Two derived variants—AMBN-249(S) and SERPINA1-341(T)—were consistently shared by *U. deningeri* and *U. spelaeus*, providing robust molecular markers for the speloid lineage. The occasional absence of these markers likely reflects limited peptide recovery rather than genuine biological variation. Their uniform presence across geographically distant sites highlights a remarkable molecular homogeneity in *U. deningeri* proteomes, in contrast with the morphological variability often described for this taxon [48].

Some Middle Pleistocene sites, however, capture intermediate stages. At Aroeira and at level TD8 of the Gran Dolina site, two individuals show both ancestral and derived AMBN-249 states, indicating incomplete fixation or population-level heterogeneity. Alternatively, this pattern may reflect residual gene flow between arctoid and speloid populations during the early stages of their divergence [14]. These findings suggest that TD8 represents a transitional horizon, bridging *U. dolinensis* from TD4 with later *U. deningeri* forms documented at TD10. The chronology of TD8, which overlaps with the Brunhes–Matuyama reversal and major climatic fluctuations (∼0.78 Ma) [49], further underscores its biochronological value to reflect both faunal turnover and ecological restructuring [50-52]. The Aroeira specimen, although inconclusive, offers a useful contrast with other sites of similar age and highlights regional variability in the pace of molecular change. While the *U. deningeri* proteome appears generally homogeneous, some populations retain ancestral residues, whereas others show early fixation of derived variants. This pattern suggests that molecular transitions within the speloid lineage were not strictly synchronous across populations. Instead, different groups likely fixed derived variants at other times, shaped by ecological or demographic pressures. Such diachronic patterns do not contradict an anagenetic model but rather indicate that cave bear evolution was gradual, though not synchronous, with populations evolving at different rates across regions.

Additional signals of variability were detected in isolated contexts. At Sala Fantasma, one specimen carried a unique AMBN-278 substitution not seen elsewhere, while others from nearby Cueva Fantasma displayed the typical speloid signature. At Monte Cucco, two individuals identified morphologically as *U. arctos* yielded proteomic profiles that did not correspond to our *U. arctos* reference, suggesting either misidentification or cryptic diversity. Finally, the specimen from Pirro Nord (Italy), attributed to *U. etruscus*, was the most degraded in the dataset. Nevertheless, the recovered segments were highly conserved, indistinguishable from both extant and extinct bear species.

Together, these results highlight the capacity of enamel proteomics to track evolutionary dynamics across temporal and geographic scales. The persistence of stable molecular signatures, combined with evidence for gradual transitions in key horizons, provides new support for an anagenetic model of cave bear evolution. By integrating molecular markers spanning more than a million years, this study opens new perspectives for molecular palaeontology and underscores the potential of enamel proteomes as tools for systematics, biochronology, and evolutionary inference.

## Materials and Methods

### Sample Selection

We analyzed 55 fossil enamel specimens from several Pleistocene sites in southwestern Europe, spanning a broad chronological and geographic range. The dataset includes individuals attributed to *U. dolinensis, U. deningeri, U. spelaeus, U. etruscus*, and *U. arctos*. Particular emphasis was placed on specimens from the Sierra de Atapuerca (Burgos, Spain)—notably Gran Dolina, Sima de los Huesos, and Cueva Fantasma—which provide key taxonomic and temporal contexts from the Early to Late Pleistocene. Complete specimen metadata are provided in ***Dataset S1 – Methodological Databases. Table S1***.

### Peptide Extraction and Identification

Enamel peptides were extracted using a digestion-free acid demineralization protocol under clean laboratory conditions at CENIEH (Burgos) and IBE (Barcelona), following previously established methods. LC-MS/MS analyses were conducted at the CRG-UPF Proteomics Core Facility (Barcelona). Protein identification used a custom reference database of ursid enamel and plasma proteins, supplemented with common contaminants and human keratins. Two search engines were employed with unspecific cleavage rules to accommodate diagenetic variation. Only proteins supported by two or more non-overlapping unique peptides and identified at <1% FDR were retained. Full analytical details are provided in ***SI. Materials and Methodology*** and ***Dataset S1 – Methodological Databases. Table S2***.

### Comparative and Phylogenetic Analysis

Ancient proteomes were compared against two reference panels: a broad Carnivora dataset and a restricted Ursidae dataset, with *Canis lupus familiaris* and *Ailuropoda melanoleuca* as outgroups. Protein sequences were aligned, treating leucine and isoleucine as isobaric. Phylogenetic inference was based on a concatenated alignment of 12 enamel-associated proteins and conducted using Bayesian and Maximum Likelihood frameworks with models fitted to each protein partition. Full parameters and dataset composition are provided in ***SI. Materials and Methodology*** and ***Dataset S1 – Methodological Databases. Table S3-4***.

## Supporting information

SI Supplementary Results

Dataset S1- Methodological Databases

SI Dataset S2- Peptide Recovery and Identification

## Data Availability

The mass spectrometry proteomics data have been deposited to the ProteomeXchange Consortium via the PRIDE [53] partner repository with the dataset identifier PXD069138.

## References

1. A. V. Sher, et al., The first record of “spelaeoid” bears in Arctic Siberia. Quaternary Science Reviews 30, 2238–2249 (2011).

2. E. Thenius, Ursidenphylogenese und Biostratigraphie (Urban und Fischer, 1959).

3. B. Kurtén, Pleistocene mammals of Europe (Routledge, 1968).

4. B. Kurtén, The cave bear story: life and death of a vanished animal. New York. (1976).

5. M. Knapp, et al., First DNA sequences from Asian cave bear fossils reveal deep divergences and complex phylogeographic patterns. Molecular Ecology 18, 1225–1238 (2009).

6. J. Krause, et al., Mitochondrial genomes reveal an explosive radiation of extinct and extant bears near the Miocene-Pliocene boundary. BMC Evol Biol 8, 220 (2008).

7. M. Pacher, A. J. Stuart, Extinction chronology and palaeobiology of the cave bear (Ursus spelaeus). Boreas 38, 189–206 (2009).

8. M. Stiller, et al., Withering away—25,000 years of genetic decline preceded cave bear extinction. Molecular Biology and Evolution 27, 975–978 (2010).

9. A. Grandal-D’Anglade, J. R. Vidal Romaní, A population study on the cave bear (Ursus spelaeus Ros.-Hein.) from Cova Eirós (Triacastela, Galicia, Spain). Geobios 30, 723–731 (1997).

10. A. Argant, Biochronologie et grands mammifères au Pléistocène moyen et supérieur en Europe occidentale : l’apport des canidés, des ursidés et des carnivores en général. Quaternaire. Revue de l’Association française pour l’étude du Quaternaire 467–480 (2009). 10.4000/quaternaire.5334.

11. G. F. Baryshnikov, A. Yu. Puzachenko, Craniometrical variability in the cave bears (Carnivora, Ursidae): Multivariate comparative analysis. Quaternary International 245, 350–368 (2011).

12. E. Santos, A. Gómez-Olivencia, M. Arlegi, J. L. Arsuaga, Cranial morphological differences within U. deningeri – U. spelaeus lineage: A double traditional and geometric morphometrics approach. Quaternary International 433, 347–362 (2017).

13. J. Wagner, S. Cermak, Revision of the early Middle Pleistocene bears (Ursidae, Mammalia) of Central Europe, with special respect to possible co-occurrence of spelaeoid and arctoid lineages. Bulletin of Geosciences 87, 461–496 (2012).

14. A. Barlow, et al., Partial genomic survival of cave bears in living brown bears. Nat Ecol Evol 2, 1563–1570 (2018).

15. M. Knapp, From a molecule’s perspective – contributions of ancient DNA research to understanding cave bear biology. Historical Biology 31, 442–447 (2019).

16. A. Barlow, et al., Middle Pleistocene genome calibrates a revised evolutionary history of extinct cave bears. Current Biology 31, 1771–1779 (2021).

17. C. Valdiosera, et al., Typing single polymorphic nucleotides in mitochondrial DNA as a way to access Middle Pleistocene DNA. Biology Letters 2, 601–603 (2006).

18. J. Dabney, et al., Complete mitochondrial genome sequence of a Middle Pleistocene cave bear reconstructed from ultrashort DNA fragments. Proceedings of the National Academy of Sciences 110, 15758–15763 (2013).

19. A. Pantoja-Pérez, J.-L. Arsuaga, The Cranium I: Neurocranium. Anat Rec (Hoboken) 307, 2278–2324 (2024).

20. N. García, J. L. Arsuaga, Les carnivores (Mammalia) des sites du Pléistocène ancien et moyen d’Atapuerca (Espagne). L’Anthropologie 105, 83–93 (2001).

21. G. Rabeder, M. Pacher, G. Withalm, Early Pleistocene Bear Remains from Deutsch-Altenburg (Lower Austria) (ÖAW, Verlag der Österreichischen Akademie der Wissenschaften, 2010).

22. J. Wagner, Pliocene to early Middle Pleistocene ursine bears in Europe: A taxonomic overview. Journal of the National Museum (Prague), Natural History Series 179, 197–215 (2010).

23. B. Demarchi, et al., Protein sequences bound to mineral surfaces persist into deep time. eLife 5, e17092 (2016).

24. R. S. Paterson, et al., Phylogenetically informative proteins from an Early Miocene rhinocerotid. Nature 643, 719–724 (2025).

25. A. J. Taurozzi, et al., Deep-time phylogenetic inference by paleoproteomic analysis of dental enamel. Nat Protoc 19, 2085–2116 (2024).

26. E. Cappellini, et al., Early Pleistocene enamel proteome from Dmanisi resolves Stephanorhinus phylogeny. Nature 574, 103–107 (2019).

27. F. Welker, et al., The dental proteome of Homo antecessor. Nature 580, 235–238 (2020).

28. F. Welker, et al., Enamel proteome shows that Gigantopithecus was an early diverging pongine. Nature 576, 262–265 (2019).

29. P. P. Madupe, et al., Enamel proteins reveal biological sex and genetic variability in southern African Paranthropus. Science 388, 969–973 (2025).

30. F. Delsuc, B. Gasse, J.-Y. Sire, Evolutionary analysis of selective constraints identifies ameloblastin (AMBN) as a potential candidate for amelogenesis imperfecta. BMC Evol Biol 15, 148 (2015).

31. D. R. Green, et al., Mapping the tooth enamel proteome and amelogenin phosphorylation onto mineralizing porcine tooth crowns. Front. Physiol. 10 (2019).

32. K. Gelse, E. Pöschl, T. Aigner, Collagens—structure, function, and biosynthesis. Advanced Drug Delivery Reviews 55, 1531–1546 (2003).

33. A. García-Vázquez, A. C. Pinto-Llona, J. Maroto, T. Torres, A. Grandal-D’Anglade, Characterising the cave bear Ursus spelaeus Rosenmüller by ZooMS: a review of peptide mass fingerprinting markers. Earth and Environmental Science Transactions of the Royal Society of Edinburgh 114, 83–93 (2023).

34. T. Asaka, et al., Type XVII collagen is a key player in tooth enamel formation. The American journal of pathology 174, 91–100 (2009).

35. P. Goettig, V. Magdolen, H. Brandstetter, Natural and synthetic inhibitors of kallikrein-related peptidases (KLKs). Biochimie 92, 1546–1567 (2010).

36. R. Sawafuji, et al., Proteomic profiling of archaeological human bone. Royal Society Open Science 4, 161004 (2017).

37. A.-K. U. Friedrich, et al., Comparative coagulation studies in hibernating and summer-active black bears (Ursus americanus). Thromb Res 158, 16–18 (2017).

38. B. A. Chow, S. W. Donahue, M. R. Vaughan, B. McConkey, M. M. Vijayan, Serum immune-related proteins are differentially expressed during hibernation in the American black bear. PLOS ONE 8, e66119 (2013).

39. K. G. Welinder, et al., Biochemical foundations of health and energy conservation in hibernating free-ranging subadult brown bear Ursus arctos. Journal of Biological Chemistry 291, 22509–22523 (2016).

40. D. A. Mugahid, et al., Proteomic and transcriptomic changes in hibernating grizzly bears reveal metabolic and signaling pathways that protect against muscle atrophy. Sci Rep 9, 19976 (2019).

41. N. Takamatsu, et al., Expression of multiple α1-antitrypsin-like genes in hibernating species of the squirrel family. Gene 204, 127–132 (1997).

42. N. Kondo, J. Kondo, Identification of novel blood proteins specific for mammalian hibernation. Journal of Biological Chemistry 267, 473–478 (1992).

43. S. Cooper, et al., Platelet proteome dynamics in hibernating 13-lined ground squirrels. Physiological Genomics 53, 473–485 (2021).

44. A. Gil-Bona, F. B. Bidlack, Tooth enamel and its dynamic protein matrix. International Journal of Molecular Sciences 21, 4458 (2020).

45. C. Froment, et al., Protein sequence comparison of human and non-human primate tooth proteomes. Journal of Proteomics 231, 104045 (2021).

46. J. P. Noonan, et al., Genomic sequencing of Pleistocene cave bears. Science 309, 597–599 (2005).

47. R. Fong-Zazueta, et al., Phylogenetic signal in primate tooth enamel proteins and its relevance for paleoproteomics. Genome Biol Evol 17, evaf007 (2025).

48. A. H. van Heteren, M. Arlegi, E. Santos, J.-L. Arsuaga, A. Gómez-Olivencia, Cranial and mandibular morphology of Middle Pleistocene cave bears (Ursus deningeri): implications for diet and evolution. Historical Biology 31, 485–499 (2019).

49. J. D. Hays, J. Imbrie, N. J. Shackleton, Variations in the Earth’s Orbit: Pacemaker of the Ice Ages. Science 194, 1121–1132 (1976).

50. J. Rodríguez, et al., One million years of cultural evolution in a stable environment at Atapuerca (Burgos, Spain). Quaternary Science Reviews 30, 1396–1412 (2011).

51. G. Cuenca-Bescós, et al., Comparing two different Early Pleistocene microfaunal sequences from the caves of Atapuerca, Sima del Elefante and Gran Dolina (Spain): Biochronological implications and significance of the Jaramillo subchron. Quaternary International 389, 148–158 (2015).

52. J. van der Made, J. Rosell, R. Blasco, Faunas from Atapuerca at the Early–Middle Pleistocene limit: The ungulates from level TD8 in the context of climatic change. Quaternary International 433, 296–346 (2017).

53. Y. Perez-Riverol, et al., The PRIDE database at 20 years: 2025 update. Nucleic Acids Res 53, D543–D553 (2025).

