## Supplementary material for "Palaeoproteomic insights into the deep roots of the cave bear lineage in Europe": SI Supplementary Results

#### SI Contents

##### I. Materials and Methodology


- I.1. Sites of Sample Provenance
- I.2. Enamel Sampling Protocol
- I.3. Protein Extraction and LC-MS/MS Analysis
- I.4. Database Construction
- I.5. Protein Identification Parameters and Search Engines
- I.6. Sequence Validation and SAP Identification

- I.7. Phylogenetic Methods and Tree Reconstruction
- I.8. Deamidation and Oxidation rates


#### II. Results


- II.1. Proteomic Recovery Intensity
- II.2. SAP Validation: Alignment Evidence and Peptide Spectra
- II.3. Expanded Phylogenetic Trees and Node Support
- II.4. Sex Determination: Peptides from AMELX/AMELY


#### III. Supplementary Dataset S1 (Excel Sheet) – Methodological Databases


- Table S1. Sample Metadata
- Table S2. Protein Databases Used for Protein Identification
- Table S3. Protein Sequences for Comparative Alignment ("Orden_Carnivora" Database)
- Table S4. Ursid-Specific Protein Database for Intra-Family Phylogenetic Analysis ("Family_Ursidae" Database)


#### IV. Supplementary Dataset S2 (Excel Sheet)– Peptide Recovery and Identification


- Table S5. Peptide Recovery Results from All Samples by Site


#### SI VI. Supplementary References

### SI I. Supplementary Materials and Methodology

#### I.1. Sites of Sample Provenance

Samples were obtained from several localities in the Iberian (Spain and Portugal) and the Italian Peninsula, representing distinct biochronological intervals of the Pleistocene. Full details are provided in ***Table S1. Sample Metadata***.

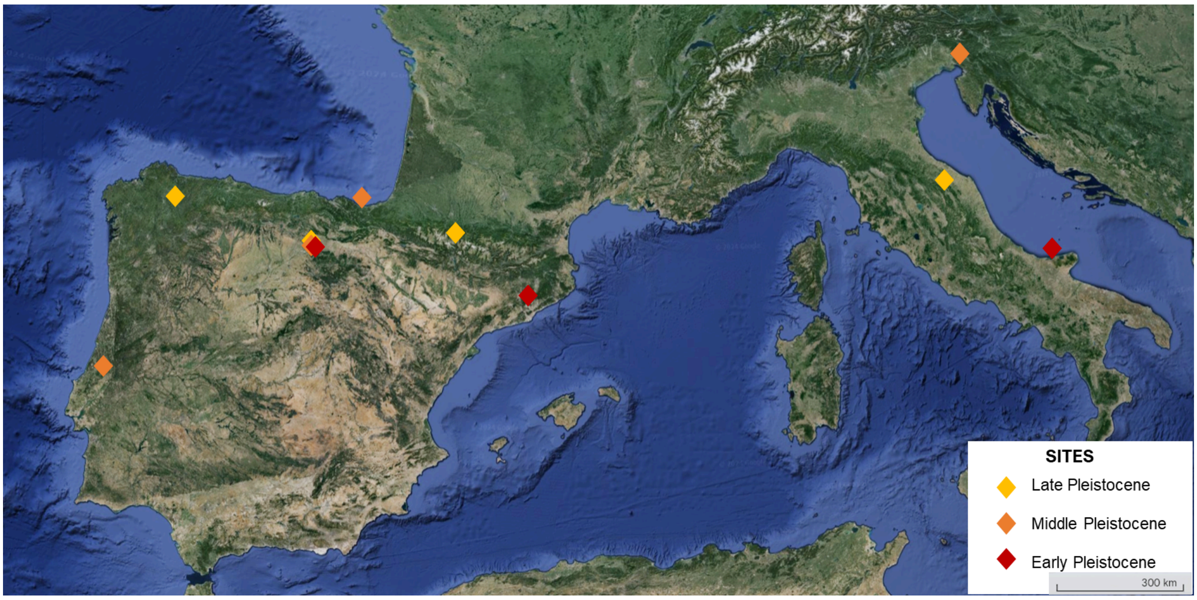


**Figure S1.** Studied sites in Spain, Portugal, and Italy: Sierra de Atapuerca (Gran Dolina, Sima de los Huesos, Fantasma), Cova Eirós, Estació Vallparadís, Cueva de Baio, Gruta de Aroeira, Monte Cucco, Pirro Nord, Visogliano. Focus on Atapuerca (Gran Dolina, Sima de los Huesos) due to its abundant and well-characterized cave bear record.

#### I.2. Enamel Sampling Protocol

To balance sample preservation and analytical yield, we adopted a set of standardized criteria aimed at minimizing damage to the specimens while maximizing protein recovery.

Enamel thickness and morphology have been extensively characterized in both extant and extinct ursids [1-3]. Cave bears exhibit notably thicker enamel than other bear species, an adaptation likely linked to their predominantly herbivorous diet [4]. In *U. spelaeus*, anterior teeth—particularly the first and second molars—display significantly greater enamel thickness compared to posterior dentition [4]. These morphological traits make molars the most suitable candidates for enamel sampling in proteomic analyses.

Whenever possible, naturally fractured specimens were prioritized to avoid damaging complete teeth. When available, enamel was sampled from existing fragments rather than from intact dental elements. For specimens excavated at Atapuerca, micro-computed tomography (micro-CT) scans were performed at the CENIEH-ICTS Microtomography and Microscopy Laboratory. All specimens were photographed before sampling to ensure full documentation of their original state. For samples sourced from external institutions, all contextual data and photographic records were meticulously compiled.

Sampling from the Atapuerca assemblage was carried out at the Geology Laboratory of the Centro Nacional de Investigación sobre la Evolución Humana (CENIEH). All manipulations took place under a laminar flow hood to maintain sterile conditions. Following separation, enamel fragments were weighed precisely and stored in sterile tubes for subsequent pulverization. Specimens held at other institutions were processed on-site.

Before extraction, any surface adhesives or glues were removed to prevent contamination or analytical interference. Cleaning was performed using a 10% sodium hypochlorite solution and disposable materials. Enamel separation was conducted using a #15 surgical scalpel blade, exploiting natural cracks when present. For unfragmented specimens, a handheld Dremel tool with a round cutting disc was employed. To minimize the inclusion of dentin peptides, residual dentin was removed using a dental bur mounted on an electric handpiece.

In total, 55 enamel samples were analyzed, each representing a unique individual. Of these, 53 originated from permanent dentition and 2 from deciduous teeth. Sampling was consistently limited to the first or second molar. The mass of enamel recovered per sample ranged from 20 to 120 mg, depending on specimen preservation and anatomical availability.

##### I.3. Protein Extraction and LC-MS/MS Analysis

All laboratory procedures, from enamel pulverization to peptide extraction, were performed in an ancient biomolecule facility at the Institut de Biologia Evolutiva (IBE, UPF-CSIC), equipped with cleanrooms operating under filtered ventilation and positive pressure. Stringent decontamination protocols were implemented, including overnight UV irradiation of surfaces, regular cleaning with 5% sodium hypochlorite followed by 70% ethanol, and the use of full protective gear (facial masks, nitrile gloves, hairnets, and disposable coveralls) to minimize the risk of cross-contamination with modern proteins. A blank extraction control was included for every set of five to seven samples. These negative controls underwent the same procedure as the enamel samples, including all reagents and consumables from the same batch, but without any biological material.

Each sample was mechanically pulverized using a granite mortar and stored in Eppendorf tubes until further processing. Demineralization was carried out using 1 mL of 10% trifluoroacetic acid (TFA) in 1.5 mL Protein LoBind® tubes. Samples were incubated at 4 °C with continuous agitation for 24 hours, a process repeated over two consecutive days to maximize peptide recovery. Each cycle included centrifugation, collection of the supernatant, and re-suspension of any residual pellet in fresh 10% TFA.

Peptide desalting and purification were performed using Stage Tips [5], prepared in-house by inserting two pre-cut 3M Empore™ C18 discs into P200 pipette tips. The tips were sequentially conditioned with methanol, 80% acetonitrile containing 0.1% TFA, and equilibrated with 0.1% TFA in water. Acidified peptides were then loaded onto the tips and retained by centrifugation (1300 × g). Bound peptides were washed with 0.1% TFA and stored at –20 °C until analysis. Elution was performed using 300 µL of 5% formic acid in 50% acetonitrile. Each sample was assigned a unique internal ID.

Mass spectrometry was conducted at the Proteomics Unit of the Centre for Genomic Regulation (CRG-UPF, Barcelona). Samples were analyzed on an Orbitrap Eclipse mass spectrometer (Thermo Fisher Scientific, San Jose, USA) coupled to an EASY-nLC 1200 nanoflow liquid chromatography system. Peptides were separated by reverse-phase LC on a 50 cm × 75 µm column packed with 2 µm C18 particles (Thermo Fisher Scientific). Chromatographic separation was achieved using a linear gradient from 5% to 25% buffer B (0.1% FA in 80% acetonitrile) over 105 minutes, followed by an increase to 40% over 15 minutes, and a final wash with 100% buffer B for 10 minutes. Flow rate was maintained at 300 nL/min. Buffer A consisted of 0.1% FA in water. The mass spectrometer operated in positive ion mode with a spray voltage of 2.4 kV and source temperature of 305 °C. Data were acquired in data-dependent acquisition (DDA) mode. Full MS scans were acquired at 120,000 resolution across an m/z range of 350–1400, using an AGC target of 4e6 and automatic injection time. The most intense precursor ions (threshold >10,000) were selected for fragmentation using a Top Speed algorithm with a 60 s dynamic exclusion. Fragmentation was performed by higher-energy collisional dissociation (HCD) at a normalized collision energy of 28%, with MS/MS spectra acquired at 30,000 resolution (AGC target: 3e4; isolation window: 0.7 m/z; maximum injection time: 54 ms). To monitor potential carry-over and instrument performance, four blanks were run before and after each sample. Digested bovine serum albumin (BSA; NEB cat. #P8108S) was used as a quality control between runs. Instrument performance was continuously assessed using QCloud [6].

#### I.4. Database Construction

To support protein identification and subsequent phylogenetic analysis, three curated protein databases were generated, each tailored to a specific stage of the analytical workflow.

- **DB_Preliminary**: This initial database comprised 296 entries corresponding to 72 proteins, obtained from three extant *Ursus* species (*U. americanus*, *U. maritimus*, and *U. arctos*) and one species from the genus *Ailuropoda* (*A. melanoleuca*). All sequences were retrieved from the NCBI public repository.
- **DB_Main**: This refined version of the preliminary set excluded sequences from *Ailuropoda melanoleuca* and incorporated additional proteins, resulting in 140 entries representing 76 proteins. It also included translated protein sequences from *Ursus thibetanus*, generated from raw whole-genome sequencing reads.
- **DB_Validation**: This final, reduced database included only proteins consistently detected across multiple samples with strong peptide support. It contained 76 sequences corresponding to 21 proteins, including entries from the NCBI RefSeq database and manually translated proteins from *Ursus thibetanus*.

In addition to retrieving annotated proteins from public databases, protein sequences were also reconstructed from modern and ancient ursid genomes with available high-throughput sequencing data. This approach allowed the inclusion of underrepresented species by applying genome-based translation pipelines adapted to coverage and data quality [7]. Sequence processing and protein translation were performed on the Castilla y León Supercomputing Infrastructure (SCAYLE; https://www.scayle.es/).

For the modern genome of *Ursus thibetanus japonicus* (~30× coverage), raw sequencing data were obtained from the ENA (DRR320311) [8], derived from blood and hair of a male individual. Reads were quality-filtered and trimmed using fastP v0.20.4 [9], discarding those shorter than 30 bp. Paired-end reads were merged and mapped to the *Ursus arctos* reference genome (RefSeq GCF_023065955.2) [10] using BWA mem v0.7.17 [11]. Duplicates were removed with biobambam v2.0.35 [12] with the bammarkduplicates command. Variant calling was performed with GATK HaplotypeCaller algorithm [13]. Gene coordinates (coding sequences) were extracted from genome annotations [10]. Protein translation was carried out in silico using internally developed Python scripts [14, https://github.com/RicardoFong/primate_enamelome]. This workflow yielded nearly complete sequences for 16 proteins, including AMELX, ENAM, AMBN, AMTN, COL17A1, COL1A1, ALB, ODAM, MMP20, SERPINC1, COL2A1, AHSG, KLK11, SERPINF1, and F2.

For *Ursus spelaeus*, an ancient genome (~2.85× coverage) from Cova Eirós (Galicia, Spain) was obtained from the ENA (ERR2678619 and ERR2678620) [15]. The individual, dated to ~31 ka [16], corresponds to specimen E-VD-1838, also analyzed proteomically in this study. Given the degraded nature of ancient DNA, a modified pipeline was used: reads were trimmed and merged using fastp, then mapped to the *Ursus arctos* genome with BWA v0.7.74 [17] with the aln algorithm. Duplicates were removed with biobambam as described before. Base calling was performed using ANGSD with dofasta option[18], coding sequences were obtained through in-house bash scripts, and protein translation followed a modified version of the previously described Python-based protocol. Five enamel-related proteins (AMBN, AMTN, MMP20, ODAM, SERPINC1) were successfully reconstructed. Missing regions were completed using consensus sequences generated with EMBOSS Cons [19] from homologous alignments within the reference set. These translated *U. spelaeus* proteins were not used for peptide identification to avoid circularity, but rather served to independently validate the paleoproteomic results from the same archaeological site, enabling direct comparison between DNA- and protein-derived sequences.

To enable sex determination based on amelogenin peptides, both AMELX and AMELY isoforms were reconstructed and included in the reference databases. Accurate, complete sequences of these isoforms are essential for distinguishing male and female individuals via peptide-based detection.

Initial searches in UniProt and NCBI revealed incomplete and low-confidence AMELX and AMELY sequences for *Ursus arctos*, a key reference species. Therefore, alternative reconstructions were performed using annotated nucleotide sequences from public genomic databases. The canonical

AMELX transcript of *Ailuropoda melanoleuca* (Ensembl v112, ENSAMET00000045715.1) was retrieved and translated. For *Ursus arctos*, the X and Y chromosome assemblies (RefSeq CM057520.1 and CM057521.1) were obtained from NCBI (PRJNA807323). Homologous regions corresponding to AMELX and AMELY exons were identified using BLAST+ v2.13.0+ with the following commands:

```
blastn -db Y_scaff_pbio_hap2.fasta -query ailme_amelx.fa -out ursarctos_amely.out0

blastn -db X_scaff_pbio_hap2.fasta -query ailme_amelx.fa -out ursarctos_amelx.out0
```

Exons were translated into amino acid sequences using the ExPASy Translate Tool [20] and assembled into full-length AMELX and AMELY proteins. These sequences were incorporated into the “DB_Principal” and “DB_Validación” peptide search databases used in MaxQuant and PEAKS.

To increase sequence diversity, AMELX proteins from *Ursus maritimus* and *Ailuropoda melanoleuca* were also included. While the reconstructed AMELY isoform from *U. arctos* is the only variant reliably used for identifying male-specific peptides, the absence of AMELY peptides in a sample may reflect either female origin or allelic loss. This limitation is consistent with previous reports showing that AMELY expression represents ~10% of AMELX levels [21].

#### I.5. Protein Identification Parameters and Search Engines

Mass spectrometry data were processed using two complementary proteomic platforms: MaxQuant v2.0.2 [22,23] and PEAKS Studio Xpro v10.6 [24]. Both software packages offer robust tools for peptide and protein identification in large-scale datasets, with features tailored to the analysis of degraded or modified ancient proteins. They allow the detection of post-translational modifications (PTMs) and provide visualization tools for interpreting complex spectra. A key advantage of PEAKS Studio Xpro lies in its highly efficient de novo sequencing capability, which enables peptide identification independent of reference databases—a particularly valuable feature for studying extinct species with limited genomic representation. A two-stage approach was adopted for the peptide search, with a final stage for sequence validation:

- **MQ1:** Initial search using MaxQuant against a broad taxonomic reference database ("DB_Preliminary"), aimed at identifying candidate enamel and collagen proteins and estimating coverage across sequences.

- **MQ2:** Advanced search using both MaxQuant and PEAKS, based on a refined database "DB_Main". This database included curated enamel and plasma protein sequences from *Ursus thibetanus japonicus* and *Ursus spelaeus*, reconstructed from published genomes, and supplemented with candidate SAPs predicted by PEAKS' SPIDER algorithm [25].

- **MQ3:** Final validation search using MaxQuant against a reduced "DB_Validation" database, incorporating validated SAPs and enriched peptide diversity.

Previous to sequence reconstruction, the results obtained in this third search were combined with the results from the advanced search in PEAKS to increase, in some cases, the coverage of the identified protein sequences. Any amino acid substitutions or variations detected by PEAKS' "SPIDER" algorithm were validated in the third advanced search in MaxQuant. Validated SAPs were included in the final version of the protein reconstruction.

MaxQuant Parameters: MaxQuant version v.2.0.2 was used. The raw data obtained were analyzed against each of the previously created databases individually and in consecutive order. In each search, the “contaminants” file included in the MaxQuant version used was incorporated. The parameters were adjusted for the search for ancient proteins. A non-specific digestion was selected. Carbamidomethylation (C) was set as a fixed modification, and Gln->pyro-Glu, Glu->pyro-Glu, Deamidation (NQ), Phospho (STY), and Oxidation (MP) were set as variable modifications. The minimum peptide length was set to 6 amino acids for non-specific searches, and the maximum was set to 20 amino acids. Peptide mass was limited to 4600 Da. The minimum score for modified and unmodified peptides was set to 40. The minimum delta score for modified and unmodified peptides was set to 0. Lastly, the parameters for de novo peptide searches using the integrated MaxNovo algorithm [26] and dependent peptides were enabled. All other parameters were set to default. Peptide filtering was performed using a false discovery rate (FDR) of 1% for peptides, peptide spectrum matches (PSMs), and proteins, with a minimum of one unique peptide per protein required.

PEAKS Parameters: The datasets submitted for the advanced MaxQuant search (MQ2) were also processed using the PEAKS program, employing its complete workflow and utilizing all search packages, from PEAKS de novo to PEAKS SPIDER, to detect potential single amino acid polymorphisms (SAPs) not present in the reference databases. A single search was performed for each sample. An error tolerance of 5 ppm was set for the precursor and 0.02 Da for fragmentation, respectively. The "Enzyme" parameter was set to "None," and the following variable modifications were included: Oxidation (M), Deamidation (NQ), N-term Pyro-Glu (Q), N-term Pyro-Glu (E), Hydroxylation (P), Phosphorylation (STY), and Carbamidomethylation (C). Carbamidomethylation (C) was included as a fixed modification, and up to 3 PTMs per peptide were allowed. The peptide filtering was performed using a false discovery rate (FDR) of 1% for peptides and a minimum protein score of -10 lgP ≥ 20. Additionally, a 95% threshold for ALC% in the de novo analysis was established, and the minimum number of unique peptides per protein was set to 1.

These parameters were calibrated considering the tandem mass spectrometer used at the Proteomics Unit.

#### I.6. Sequence Validation and SAP Identification

For the reconstruction of proteins from ancient peptide fragments, a combination of bioinformatic analysis techniques and manual validation was employed. Peptides identified in MaxQuant searches were aligned to reference proteins to generate sample-specific FASTA reconstructions. This process followed an adapted semi-automated workflow [27] implemented in R (https://www.R-project.org/). The script integrated three MaxQuant output files—*summary.txt*, *msms.txt*, and *evidence.txt*—and mapped peptide coordinates to the "DB_Validation" reference database. Any discrepancies in the reconstructed sequences were carefully reviewed manually, particularly in cases where protein sequences were suggested by de novo tools integrated into MaxQuant and PEAKS. A manual exclusion list was applied to filter out peptides based on the following criteria: (1) Poor ion coverage or low-quality MS/MS fragmentation spectra; (2) Low signal intensity in the mass spectrum; (3) Peptides likely of microbial origin.

Following the MaxQuant-based reconstruction, additional peptides uniquely identified by PEAKS were integrated under strict conditions:

- **Analytical consistency**: Although absent in MaxQuant, these peptides were consistently recovered by PEAKS with high-quality spectral support.
- **Relevance through PEAKS analysis:** Although these peptides did not appear in MaxQuant results, their detection in PEAKS indicated their significance and justified their inclusion. This discrepancy likely arises from the higher sensitivity and specificity of PEAKS in certain analyses.
- **Spectral quality**: Spectra with strong, well-resolved fragment ions provided sufficient evidence to support peptide inclusion, even when represented by a limited number of PSMs.
- **Presence of ancient-specific post-traductional modifications (PTMs):** Modifications such as deamidation or hydroxylation—indicative of diagenetic processes—further substantiated the endogenous origin of the peptides.

Final sequences were exported in FASTA format. Ambiguous residues that could not be confidently assigned were denoted by a hyphen ("–") to indicate gaps in the sequence reconstruction.

To validate the identification of Single Amino Acid Polymorphism (SAPs) with potential phylogenetic relevance, a multi-criteria framework was employed:

- **Redundant Peptide Spectrum Match (PSM) coverage**: Confidence in SAPs increased when supported by multiple overlapping peptides showing identical substitutions [28, 29].
- **BLAST validation**: Peptides were aligned against public protein databases to confirm taxonomic and positional consistency.
- **Fragment ion inspection**: High-resolution spectra were manually examined to assess fragment ion series across variant positions.
- **PTM consideration**: Potential interference from post-translational modifications (e.g., deamidation) was carefully evaluated.
- **Nucleotide substitution plausibility**: The number and type of nucleotide changes required for each amino acid substitution were assessed to rule out improbable variants.

The complementary use of ancient DNA and protein analyses has proven highly effective in evolutionary and phylogenetic studies, as each method can serve to validate findings from the other. The integration of multiple molecular approaches ensures robust validation and provides consistent phylogenetic insights, particularly in the case of the *U. spelaeus* samples analyzed in this study. Whenever possible, proteomic data were cross-referenced with protein sequences predicted from ancient *U. spelaeus* genomes. This integrative strategy—combining paleoproteomic and paleogenomic data—provides complementary lines of evidence for sequence variation and enhances phylogenetic resolution [30, 31]. Such convergence is particularly valuable in cases where DNA and protein preservation overlap, as demonstrated in the cave bear specimens analyzed here.

#### I.7. Phylogenetic Methods and Tree Reconstruction

The selection of proteins suitable for comparative and phylogenetic analysis was based on multiple criteria related to both protein function and sequence variability. First, enamel-specific proteins were prioritized, given their established use in previous phylogenetic studies of extinct mammals [27,29, 32]. These include proteins directly involved in enamel formation and mineralization. Additionally, other highly abundant proteins—despite not being traditionally associated with the enamel proteome—were also considered when partially reconstructed with reliable coverage. Second, the level of sequence variation in each protein was evaluated by quantifying the number of validated SAPs relative to the reference sequence. Proteins exhibiting variability across multiple samples were prioritized, as such polymorphisms provide valuable phylogenetic signals. Third, proteins showing potential heterozygous positions were also taken into consideration. Due to FASTA format constraints, only one amino acid per position can be reported. In these cases, the variant supported by the highest number of high-confidence peptide-spectrum matches (PSMs) was selected. When the derived (mutated) amino acid was already present in the protein reference, it was assumed to represent the fixed state, and thus retained in the reconstruction. This strategy ensures that the selected sequences reflect the most probable and phylogenetically informative variant at each position. Based on these criteria, the following twelve proteins were selected for final phylogenetic analysis:

AHSG, ALB, AMBN, AMELX, AMTN, COL1A1, COL17A1, ENAM, MMP20, ODAM, SERPINA1, SERPINC1

To increase sequence coverage and improve the robustness of phylogenetic inference, a strategy of intra-specific sequence merging was employed. Samples from the same species, archaeological site, and stratigraphic level were combined to generate consensus protein sequences, thereby enhancing signal in regions with low individual coverage. Sequence merging was performed using the EMBOSS Cons tool (European Bioinformatics Institute, EMBL-EBI; https://www.ebi.ac.uk/), which generates consensus sequences from multiple alignments [19].

- For *Ursus dolinensis*, all specimens from level TD4 of Gran Dolina were merged.
- For *Ursus deningeri*, samples from Sima de los Huesos, Cueva Mayor, and Sala de las Oseras were combined.

- For *Ursus spelaeus*, sequences from Cova Eirós and Coro Tracito were merged.

This strategy significantly improved overall sequence coverage and enabled inclusion of polymorphic regions that would otherwise have remained unrepresented, thus providing greater resolution for downstream phylogenetic analyses.

To contextualize the ancient protein sequences within a broader phylogenetic framework, two comparative databases were assembled: one encompassing multiple carnivoran families and a second focused on members of the family Ursidae. These databases served as references for multiple sequence alignments and phylogenetic analyses.

Carnivora Reference Database ("*Carnivora Dataset*"): This database includes 12 proteins from 11 species across six families within the Carnivora order, for a total of 125 curated entries. All sequences were retrieved from the NCBI public repository, either as fully annotated proteins or reconstructed from published genomes. Taxonomic representation is summarized as follows:

- Five species from the family Ursidae, four of which belong to the genus *Ursus* (*Ursus arctos, Ursus maritimus, Ursus americanus,* and *Ursus thibetanus*), and one from the genus *Ailuropoda* (*Ailuropoda melanoleuca*).
- One representative from the family Canidae, *Canis lupus familiaris*.
- Two representatives from the family Mustelidae, which includes weasels and relatives (*Mustela erminea* and *Mustela putorius furo*).
- Two representatives from the family Phocidae, the harbor seal (*Phoca vitulina*) and the southern elephant seal (*Mirounga leonina*).
- One representative from the family Odobenidae, the walrus (*Odobenus rosmarus*).
- One representative from the family Felidae, the tiger (*Panthera tigris*), whose sequences served as the outgroup in the analysis.

In cases where protein sequences were unavailable in the databases, they were computationally reconstructed from genomic data.

Ursid Reference Database ("*Ursidae Dataset*"): A second, taxonomically focused database was built to examine phylogenetic relationships within the family *Ursidae*. This dataset comprises 51 entries across 12 proteins, representing the following taxa:

- A single representative species from the genus *Ailuropoda* (*Ailuropoda melanoleuca*).
- Four representative species from the genus *Ursus* (*Ursus arctos, Ursus maritimus, Ursus americanus*, and *Ursus thibetanus*).
- The reference sequences of the species *Canis lupus familiaris* served as the outgroup for the analysis.

Except for the sequences from *Ursus thibetanus* and some manually reconstructed sequences, such as the AMELX protein for *Ursus arctos*, the remaining sequences were sourced from proteins published in the reference NCBI repositories. In certain cases, sequences for specific proteins were unavailable. All reconstructed ancient protein sequences and reference proteins were aligned using MAFFT v7.520 [33]. Reference proteins from the comparative database were aligned using the following parameters:

```
mafft --amino --maxiterate 1000 --globalpair reference_db.fasta > aligned_reference_db.fasta
```

Reconstructed sequences from ancient samples were added using the *--addfragments* method to minimize misalignment due to gaps or missing data:

```
mafft --reorder --keeplength --addfragments cavebear_samples.fasta --auto aligned_reference_db.fa > final_msa_cavebear.fa
```

All resulting alignments were manually inspected and corrected to resolve minor misalignments. To retain only the most phylogenetically informative regions, alignments were filtered using GBlocks [34], which removes ambiguous or poorly aligned regions. To address the ambiguity caused by isobaric amino acids leucine (L) and isoleucine (I), which are indistinguishable in ancient proteomics, all isoleucine residues were converted to leucines at variable positions. This normalization was applied consistently across all sequences to avoid artificial variation in the alignment. Signal peptides, when present, were removed based on annotations from UniProt, with the exception of *COL17A1*, where the signal peptide was retained. Final alignments for each ancient specimen were concatenated manually in the following order of 12 selected proteins:

AHSG, ALB, AMBN, AMELX, AMTN, COL1A1, COL17A1, ENAM, MMP20, ODAM, SERPINA1, SERPINC1

Phylogenetic relationships were inferred from the concatenated protein alignments using both maximum likelihood (ML) and Bayesian inference (BI) methods:

<u>ML Analyses:</u> **IQ-TREE v1.6.12** [35] was used to infer ML trees with partitioned models and branch support estimates using 5,000 replicates each of ultrafast bootstrap (UFBoot):

```
iqtree -nt 4 -s Namefile_concatenated.fasta -spp $partfile -alrt 5000 -bb 5000 -nstop 500 -nm 10000 -wsplits -pre $outdir
```

Protein-specific evolutionary models were determined using ModelFinder [36]:

```
iqtree -s Namefile_concatenated. fasta -nt {{threads}} -m MF -msub nuclear
```

Analyses were also conducted with **PhyML v3.148** [37] using the concatenated alignment, converted from FASTA to PHYLIP format. Each analysis was initiated from multiple random starting trees under the JTT substitution model, with empirical amino acid frequencies. The gamma shape parameter and the proportion of invariable sites were estimated by maximum likelihood. Node support was assessed through 1,000 bootstrap replicates. The exact command used was:

phyml-mpi -i Namefile_concatenated.phy -d aa -b 1000 -m JTT -a e -s BEST -v e -o tlr -f m --rand_start --n_rand_starts 4 --r_seed $RAND --print_site_lnl --print_trace --no_memory_check

Bayesian Inference: **MrBayes v3.2.7a** [38] was used with the same partitioning scheme as in IQ-TREE. Each protein was defined as a separate partition with unlinked parameters. The *mixed* amino acid model prior was used to allow model averaging across substitution matrices:

set autoclose=yes;

execute Namefile_concatenated.nex;

charset AHSG=1-248;

charset ALB=249-837;

charset AMBN=838-1151;

charset AMELX=1152-1326;

charset AMTN=1327-1369;

charset COL1A1=1370-2647;

charset COL17A1=2648-4129;

charset ENAM=4130-5239;

charset MMP20=5240-5700;

charset ODAM=5701-5964;

charset SERPINA1=5965-6333;

charset SERPINC1=6334-6614;

partition ConcatenationProteins = 12: AHSG, ALB, AMBN, AMELX, AMTN, COL1A1, COL17A1, ENAM, MMP20, ODAM, SERPINA1, SERPINC1;

set partition = ConcatenationProteins;

prset aamodelpr = mixed;

mcmc nchains = 8 nruns=4 ngen = 3000000 samplefreq=100 printfreq=100 diagnfreq=1000;

sump;

sumt relburnin = yes burninfrac = 0.25;

quit;

All phylogenetic trees were visualized and annotated using FigTree v1.4.4 (FigTree). Final trees display both ML bootstrap values and Bayesian posterior probabilities where applicable.

##### I.8. Deamidation and Oxidation rates

Deamidation of glutamine (Q) and asparagine (N), together with oxidation of methionine (M) and proline (P), was quantified using a custom script. The script calculates conversion ratios between modified and unmodified residues, normalized to peptide spectrum matches (PSMs), and validated by 1000 bootstrap replicates. Early Pleistocene specimens displayed more advanced diagenesis, with lower yields and shorter peptides.

### SI II. Supplementary Results

#### II.1. Proteomic Recovery Intensity

Relative peptide intensity (log10-transformed) across all samples compared with the corresponding number of peptide-spectrum matches (PSMs), Figure 2.A of the Main Text. Overall, Pleistocene specimens show comparable values of relative intensity, while the modern brown bear displays the expected abundance for a fresh enamel sample despite yielding fewer PSMs than some Middle and Late Pleistocene individuals. This discrepancy likely reflects differences in spectrum assignment rather than actual protein abundance, since intensity provides a direct measure of peptide signal whereas PSM counts depend on database matching and fragmentation efficiency. Thus, intensity serves as a complementary metric of recovery, highlighting that peptide abundance is consistent with preservation status even when PSM counts differ. All specimens were analyzed on the same mass spectrometry platform using an identical extraction and LC-MS/MS protocol.

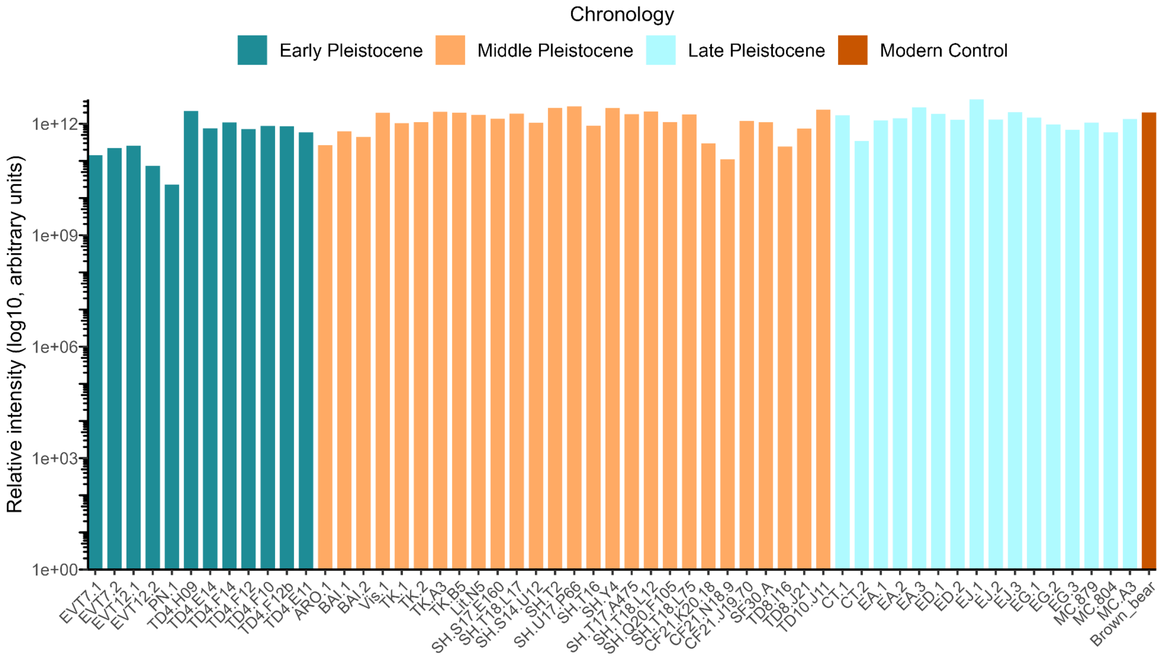


**Figure S3.** Relative peptide intensity (log10) across all enamel samples. All samples were analyzed using the same mass spectrometry platform and identical extraction and LC-MS/MS protocols.

#### II.2. SAP Validation: Alignment Evidence and Peptide Spectra

The distribution of amino acid substitutions across enamel and serum proteins highlights contrasting patterns of variation. Enamel proteins such as ameloblastin (AMBN) and enamelin (ENAM) show the highest number of SAPs, whereas others, including MMP20 and AHSG, display only a few sites of variation. Several SAPs appear uniquely in single specimens, suggesting localized or lineage-specific changes, while others are shared across multiple taxa. Figure S4 illustrates where these substitutions accumulate along the protein sequences, emphasizing regions that may be more prone to evolutionary change.

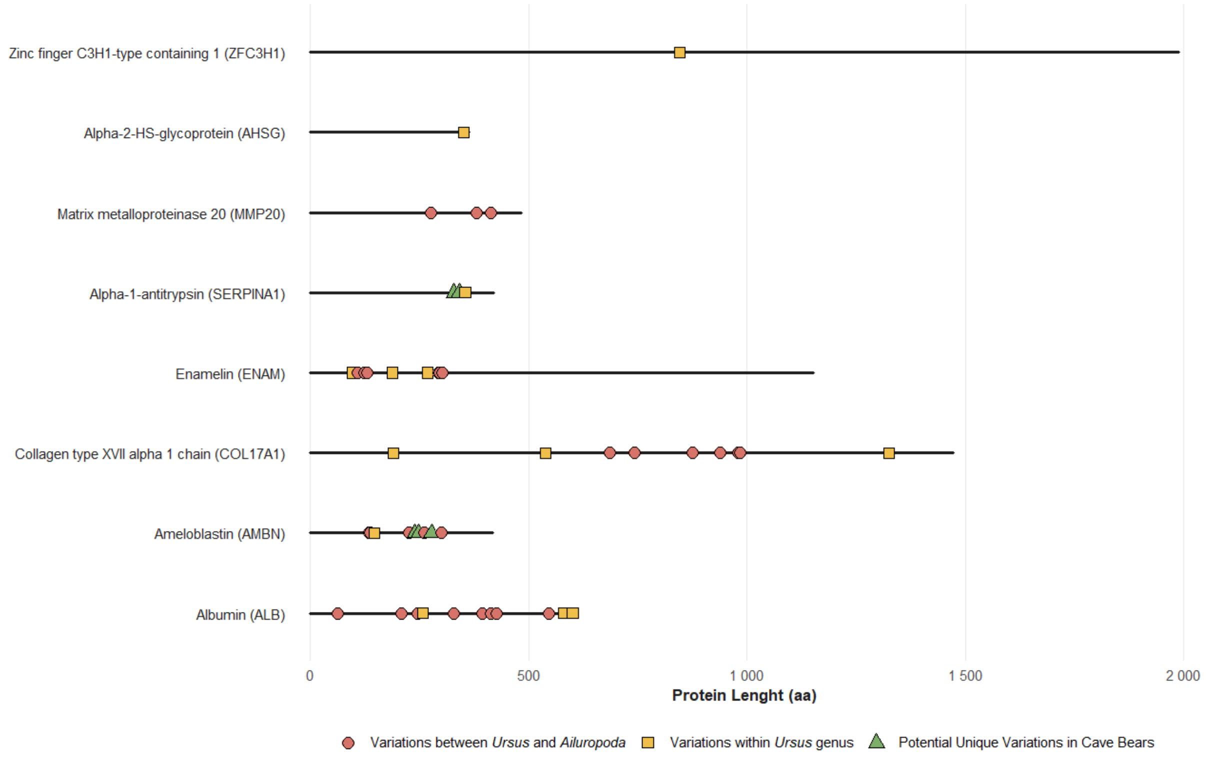


**Figure S4.** Distribution of single amino acid polymorphisms (SAPs) across protein sequences. Lollipop plot depicting variable amino acid positions along enamel and serum proteins. Each marker represents a SAP, scaled by frequency.

**SAPs and Position by Protein**

| | | | | | | | | | | | |
|---|---|---|---|---|---|---|---|---|---|---|---|
| Albumin (ALB) | 64 | 208 | 246 | 258 | 328 | 395 | 414 | 426 | 547 | 580 | 601 |
| Ameloblastin (AMBN) | 134 | 136 | 137 | 146 | 226 | 227 | 240 | 249 | 262 | 278 | 301 |
| Collagen type XVII alpha 1 chain (COL17A1) | 191 | 538 | 686 | 743 | 875 | 939 | 981 | 985 | 1,325 | | |
| Enamelin (ENAM) | 97 | 109 | 123 | 131 | 188 | 269 | 294 | 297 | 303 | | |
| Alpha-1-antitrypsin (SERPINA1) | 329 | 341 | 355 | | | | | | | | |
| Matrix metalloproteinase 20 (MMP20) | 276 | 381 | 413 | | | | | | | | |
| Alpha-2-HS-glycoprotein (AHSG) | 351 | | | | | | | | | | |
| Zinc finger C3H1-type containing 1 (ZFC3H1) | 846 | | | | | | | | | | |

■ Variations between *Ursus* and *Ailuropoda* ■ Variations within *Ursus* genus ■ Potential Unique Variations in Cave Bears

**Table S1.** Single amino acid polymorphisms (SAPs) identified in enamel and serum proteins. Summary of proteins with their respective number and positions of SAPs, including novel variants.

##### II.2.1. Validation of SAP AMBN-249

The AMBN-249 substitution [tyrosine (Y) → serine (S)] was consistently detected across all analyzed specimens of *U. deningeri* and *U. spelaeus*, as well as in several individuals classified as *Ursus* sp., including three from Cueva Fantasma (see Figure 2 in the Main Text). Representative spectral evidence of selection of samples is provided here, including annotated MS/MS spectra and fragment ion series supporting the variant residue.

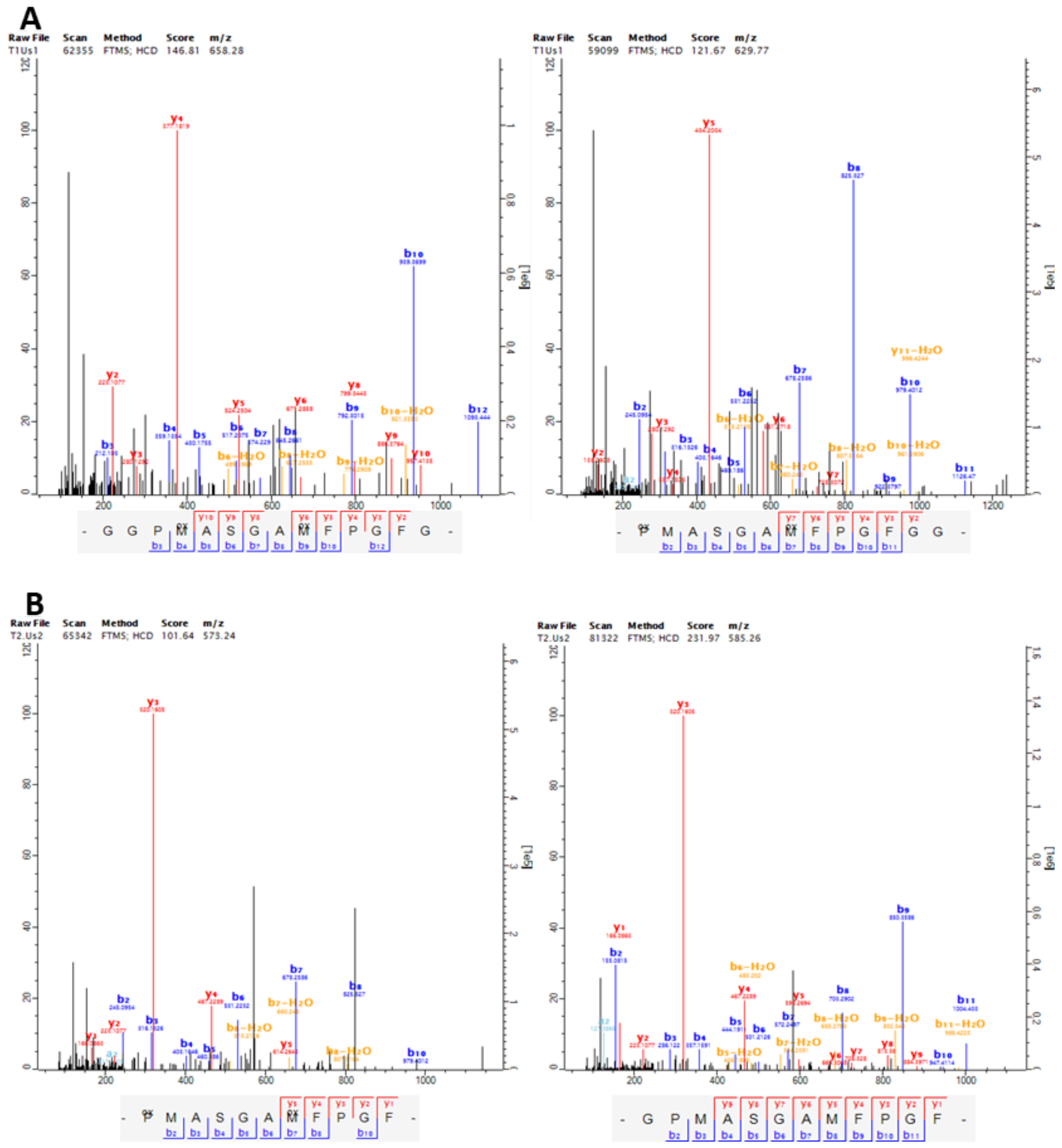


**Figure S5.** Selection of mass spectra and corresponding fragment ions covering AMBN-249 obtained with MaxQuant from *Ursus spelaeus* specimens. (A) Sample CT.1. (B) Sample CT.2.

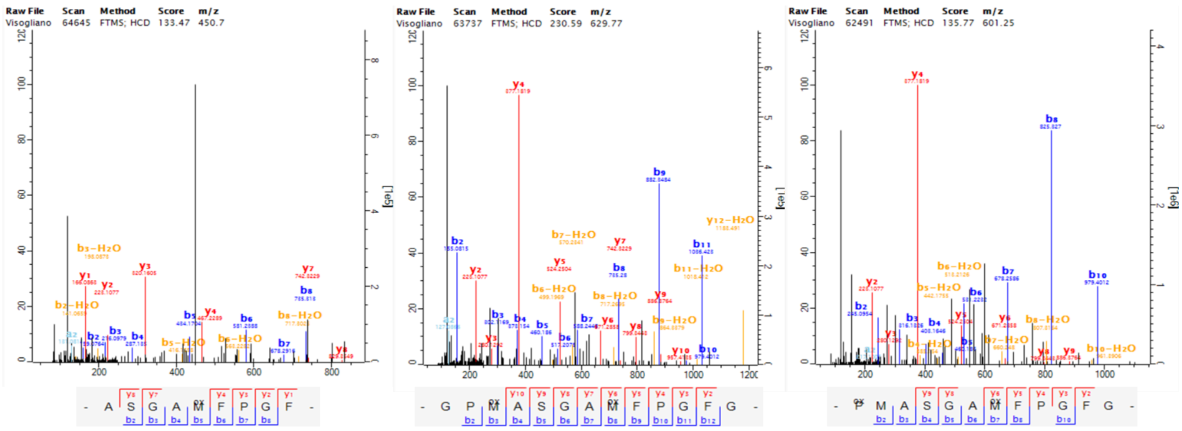


**Figure S6.** Selection of mass spectra and corresponding fragment ions covering AMBN-249 obtained with MaxQuant in Vis.1 sample *Ursus deningeri.*

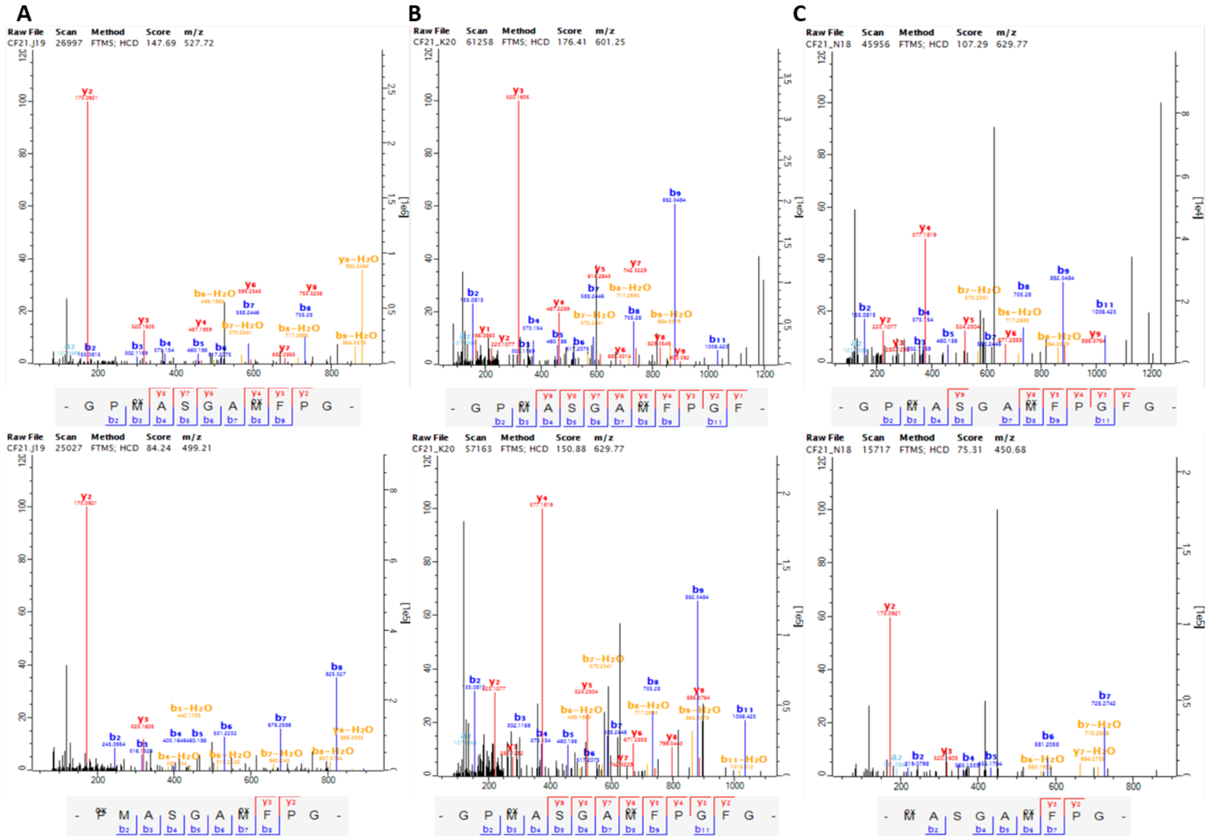


**Figure S7.** Selection of mass spectra and corresponding fragment ions covering AMBN-249 obtained with MaxQuant. (A) Unique peptides in CF21.J19. (B) Unique peptides in CF21.K20. (C) Unique peptides in CF21.N18.

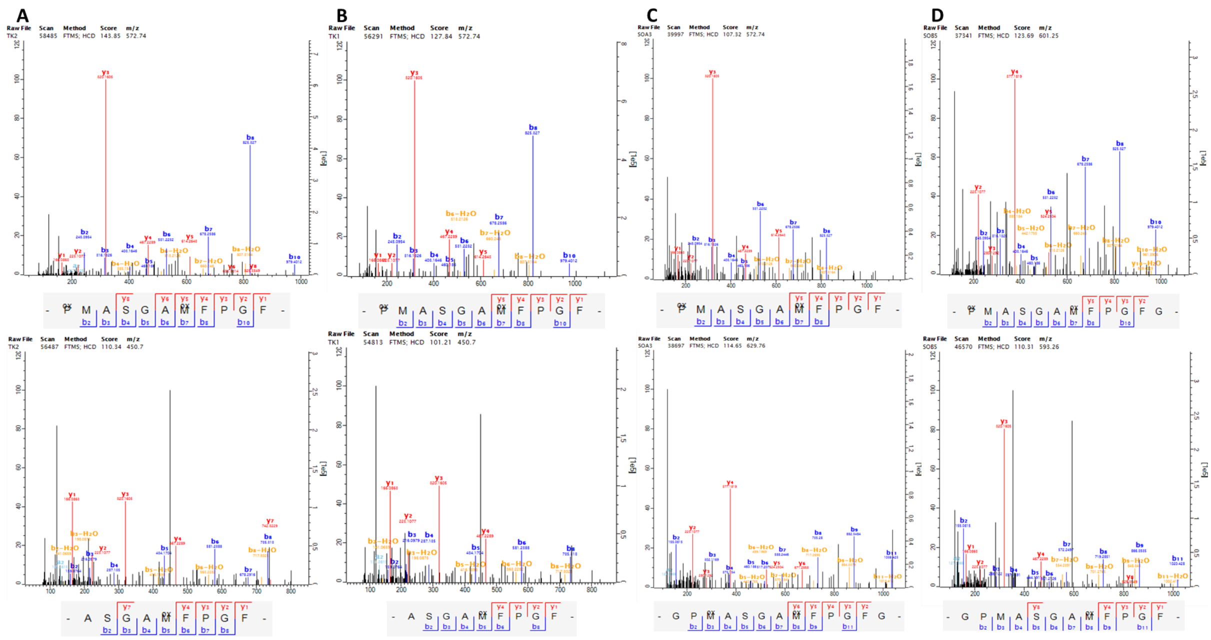

**Figure S8.** Selection of mass spectra and corresponding fragment ions covering AMBN-249 obtained with MaxQuant from *Ursus deningeri* specimens. (A) Sample TK.2. (B) Sample TK.1. (C) Sample TK.A3. (D) Sample TK.B5.

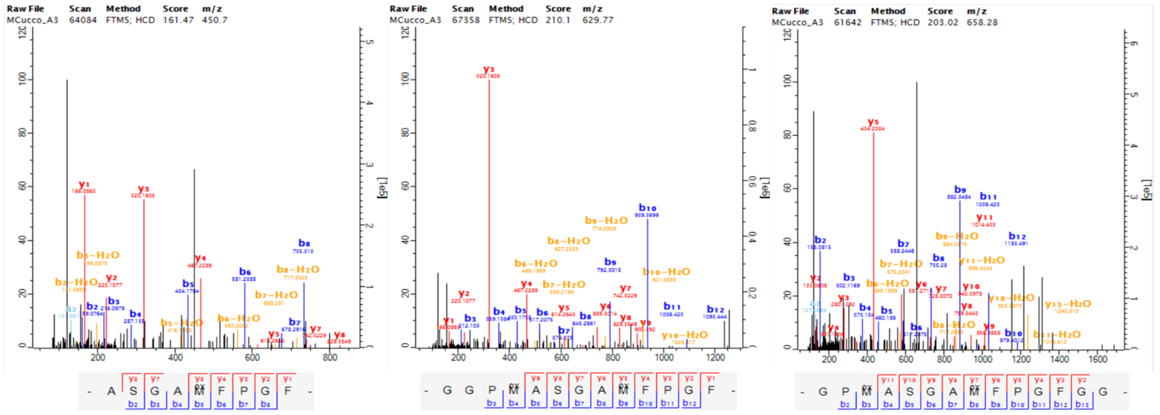

**Figure S9.** Selection of mass spectra and corresponding fragment ions covering AMBN-249 obtained with MaxQuant in MC.A3 sample *Ursus spelaeus*.

In addition, two specimens (*ARO.1* and *TD8.J21*) show evidence of heterozygosity at this position: both the reference and substituted peptide variants were identified within the same individual, with co-occurring spectra supporting true heterozygosity rather than analytical noise. All corresponding spectra are presented below, with annotated ion series confirming each variant. This amino acid replacement is notable for its widespread presence in Middle and Late Pleistocene cave bear specimens. Its recurrence across temporally and geographically distinct populations supports the hypothesis of a heritable variant that became fixed within cave bear lineages during this period.

<u>Sample TD8.J21</u> from TD8 level in Gran Dolina Site, shows clear evidence of heterozygosity, with both tyrosine (Y) and serine (S, derived state) variants detected. Independent identifications from MaxQuant and PEAKS confirmed the presence of both forms in balanced proportions, ruling out

analytical noise. Consistent post-translational modifications were observed in spectra of both variants.

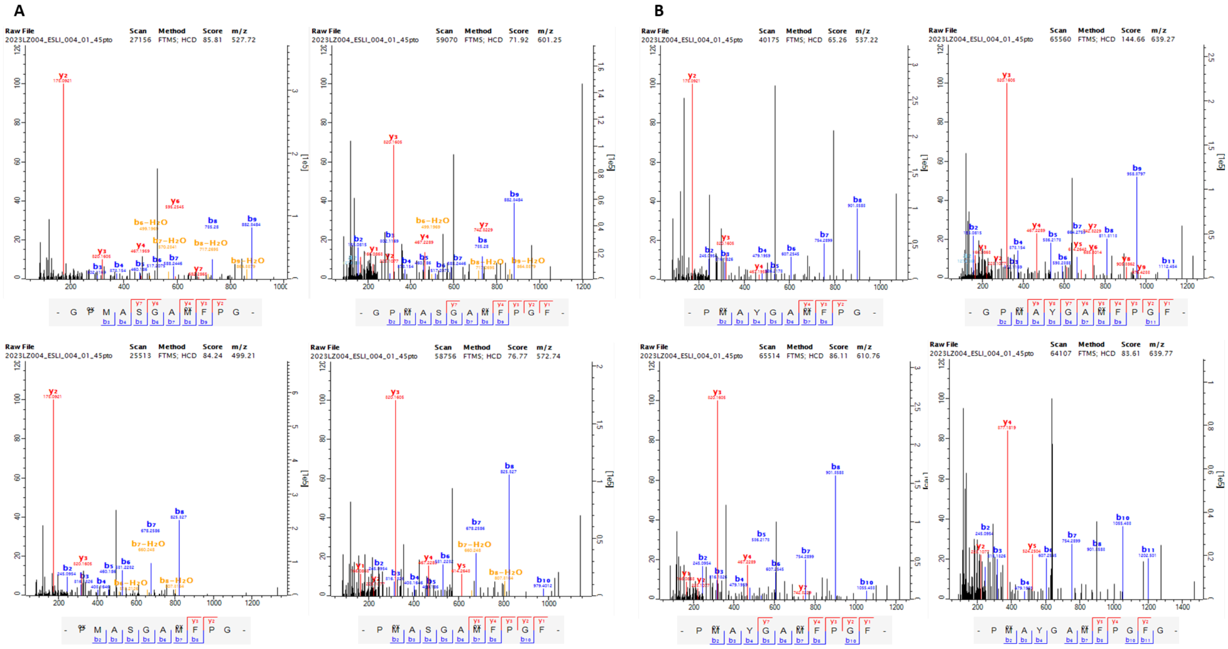


**Figure S10.** Mass spectra and corresponding fragment ions covering AMBN-249 in TD8.J21 *Ursus deningeri*, obtained with MaxQuant. (A) Peptides containing S. (B) Peptides containing Y.

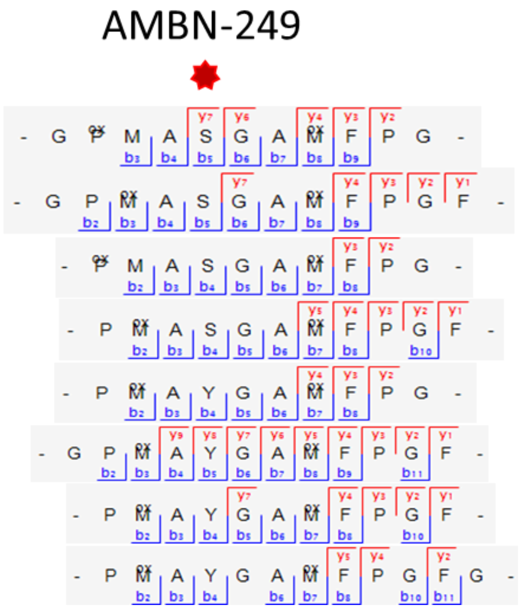


**Figure S11.** Peptide and fragment-ion coverage of AMBN; position 249 in specimen TD8.J21, marked with a red asterisk.

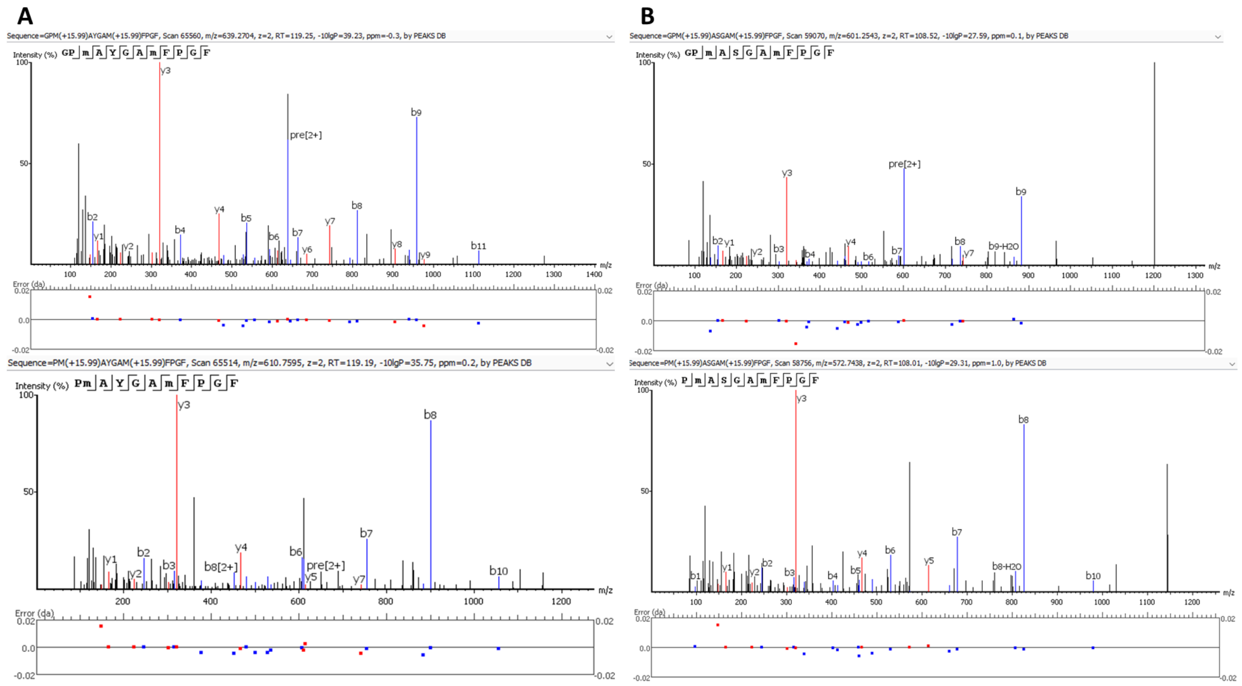


**Figure S12.** Mass spectra covering AMBN-249 in TD8.J21 *Ursus deningeri* obtained with PEAKS. (A) Unique peptides containing Y. (B) Unique peptides containing S.

<u>Sample ARO.1</u> from Aroeira Site, shows tentative evidence for heterozygosity. MaxQuant identified a peptide carrying serine (S), while PEAKS recovered both Y-containing peptides and one additional S peptide supported by a low-quality spectrum. Although engine-specific discrepancies and limited spectral support caution against over-interpretation, the co-occurrence of Y and S at this site is consistent with putative heterozygosity in this specimen.

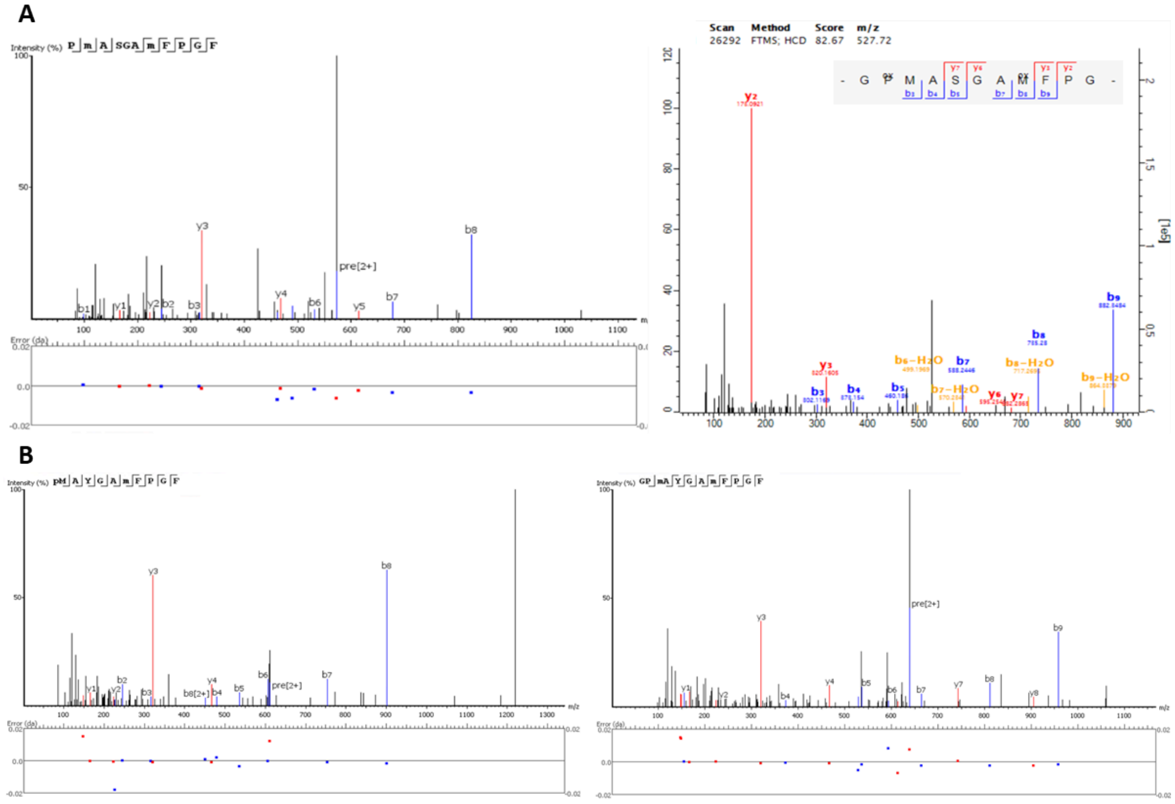


**Figure S13.** Mass spectra and corresponding fragment ions covering AMBN-249 in ARO.1 *Ursus deningeri* (A) Unique peptides containing S obtained by PEAKS and MaxQuant. (B) Unique peptides containing Y obtained by PEAKS.

##### II.2.2. Validation of SAP AMBN-278

The AMBN-278 substitution was identified exclusively in the Sala Fantasma specimen SF30.A. This variant is supported by multiple overlapping peptides with high-quality mass spectra in both search engines, confirming the presence of a residue change from leucine (L) to valine (V). The most likely explanation is a single nucleotide substitution in the codon for leucine (CUU, CUC, CUA, or CUG) into a codon for valine (GUU, GUC, GUA, or GUG), consistent with a transversion at the first nucleotide position. Both leucine and valine are hydrophobic and nonpolar, suggesting that this substitution is unlikely to drastically alter the protein’s structure or function.

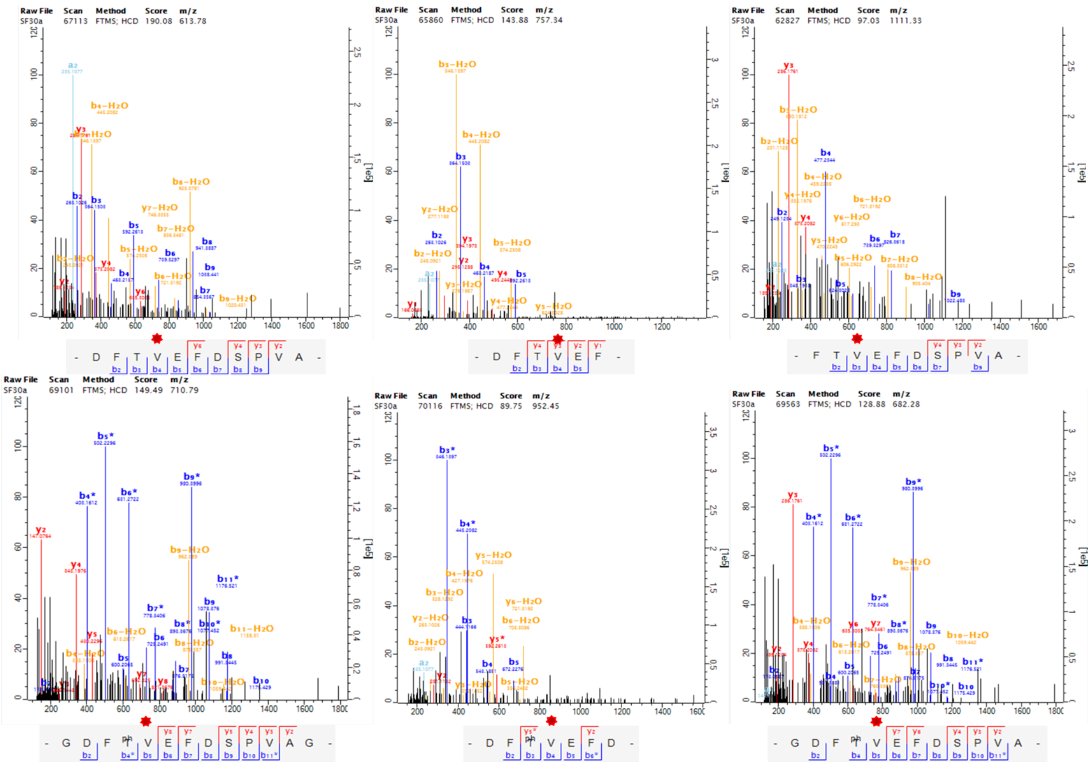

**Figure S14.** Mass spectra and corresponding fragment ions covering AMBN-278 in SF30.A *Ursus sp.* using MaxQuant, highlighted by a red asterisk.

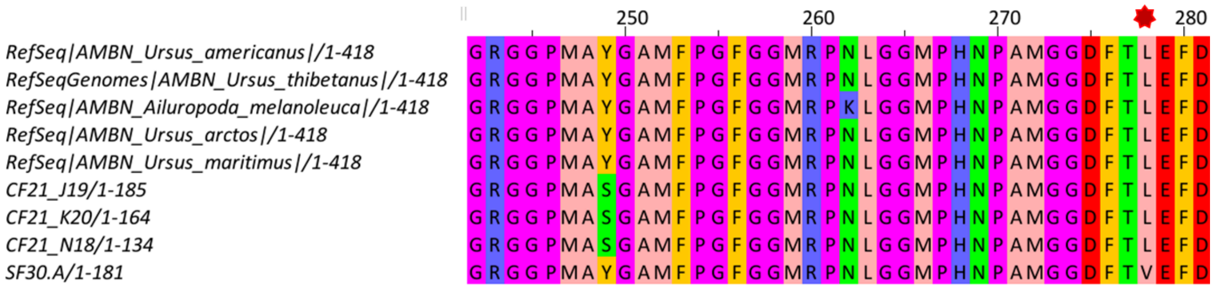


**Figure S15.** Multiple sequence alignment of the ameloblastin (AMBN) protein generated with Jalview using the comparative dataset, showing the variable positions AMBN-249 and AMBN-278, with the latter highlighted by a red asterisk.

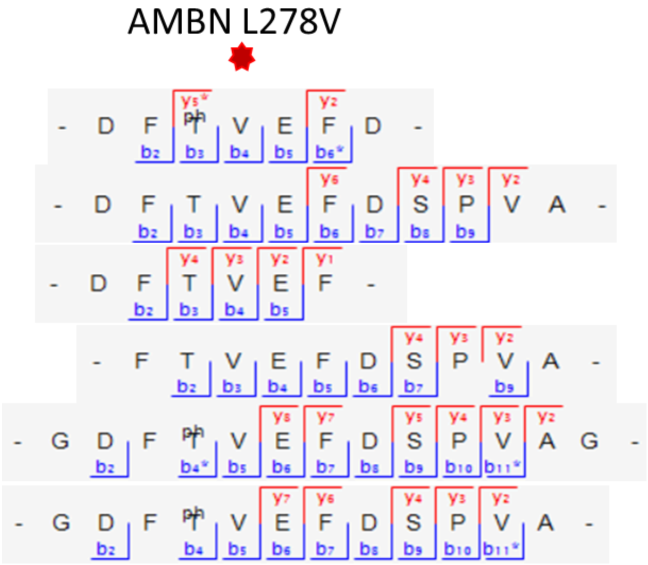


**Figure S16.** Peptide and fragment-ion coverage of AMBN; position 278 in specimen SF30.A (*Ursus* sp.) marked with a red asterisk.

##### II.2.3. Validation of SAP SERPINA1-341

The substitution at SERPINA1-341 [asparagine (N) → threonine (T)], was consistently detected across the majority of Middle and Late Pleistocene specimens analyzed. Numerous MS/MS spectra independently identified with both MaxQuant and PEAKS support the presence of this variant in specimens of *U. deningeri* and *U. spelaeus*, as well as in individuals classified as *Ursus* sp. and in one specimen of *U. arctos*. Early Pleistocene samples, in contrast, consistently retained the ancestral asparagine state. The recurrence of the variant in temporally and geographically distinct Middle and Late Pleistocene cave bears indicates a heritable mutation established within these populations

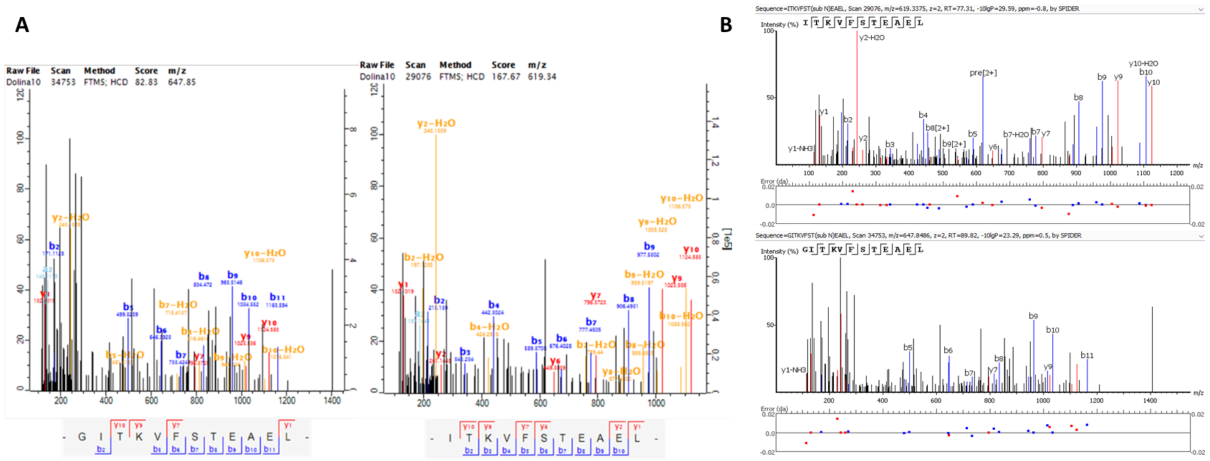


**Figure S17.** Mass spectra and corresponding fragment ions covering SERPINA1-341 in TD10.4.J11 *Ursus deningeri.* (A) Unique peptides containing Threonine obtained by MaxQuant. (B) Unique peptides containing Threonine obtained by PEAKS.

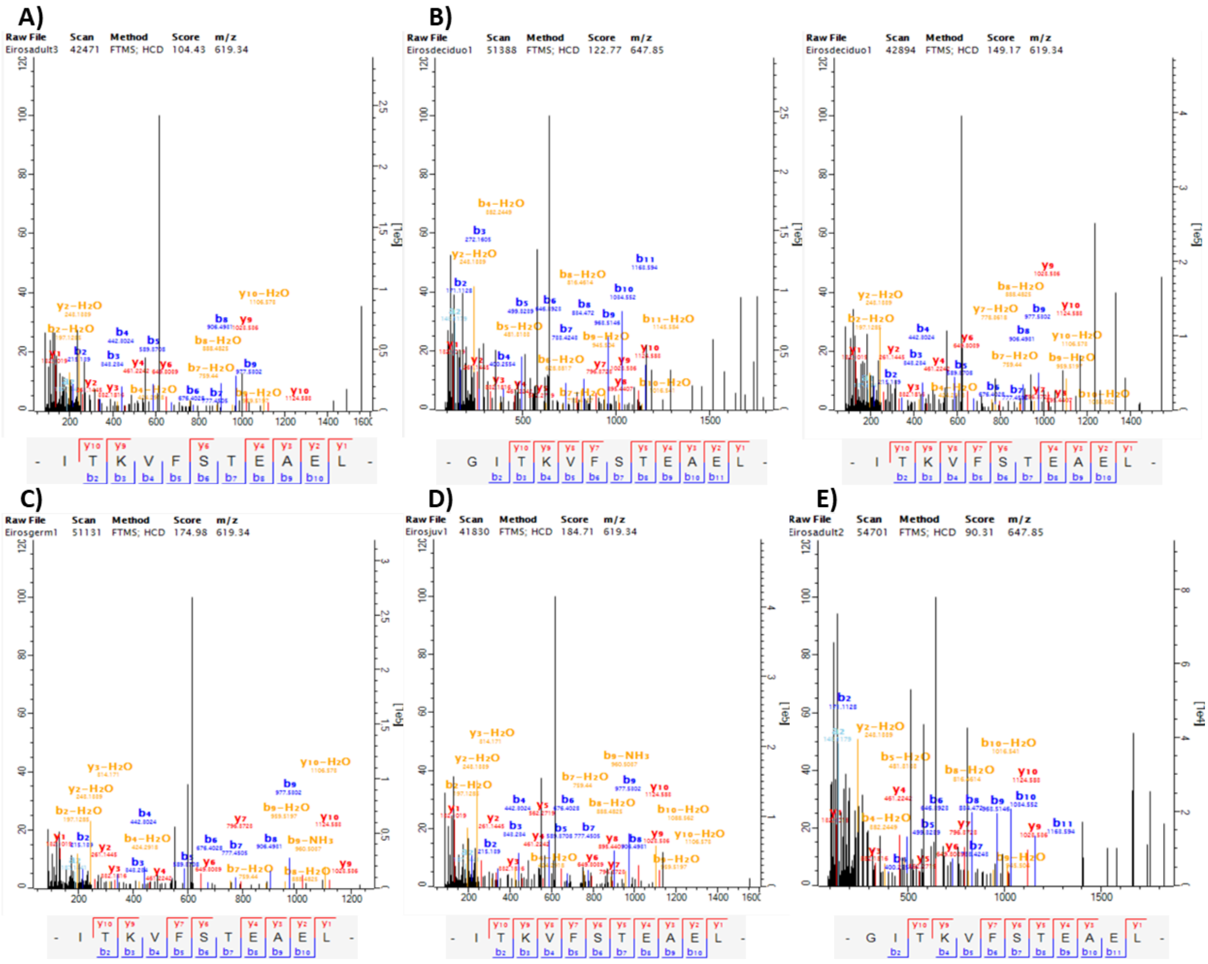

**Figure S18.** Selection of mass spectra and corresponding fragment ions covering SERPINA1-341 obtained with MaxQuant from *Ursus spelaeus* specimens. (A) Sample EA.3. (B) Sample Ed.1. (C) Sample EG.1. (D) Sample EJ.1. (E) Sample EA.2.

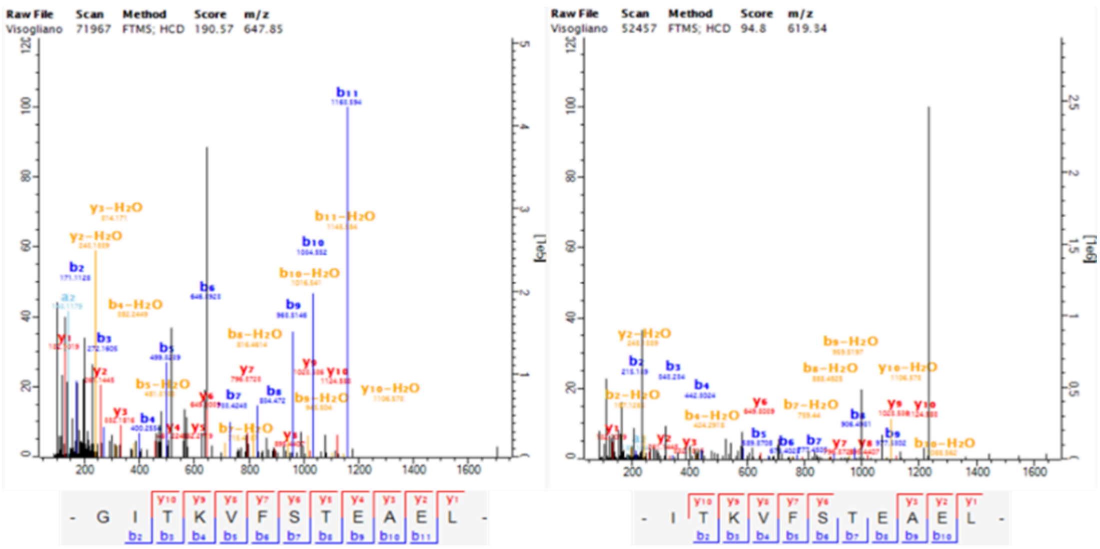

**Figure S19.** Mass spectra and corresponding fragment ions covering SERPINA1-341 in Vis.1 *Ursus deningeri* specimen using MaxQuant.

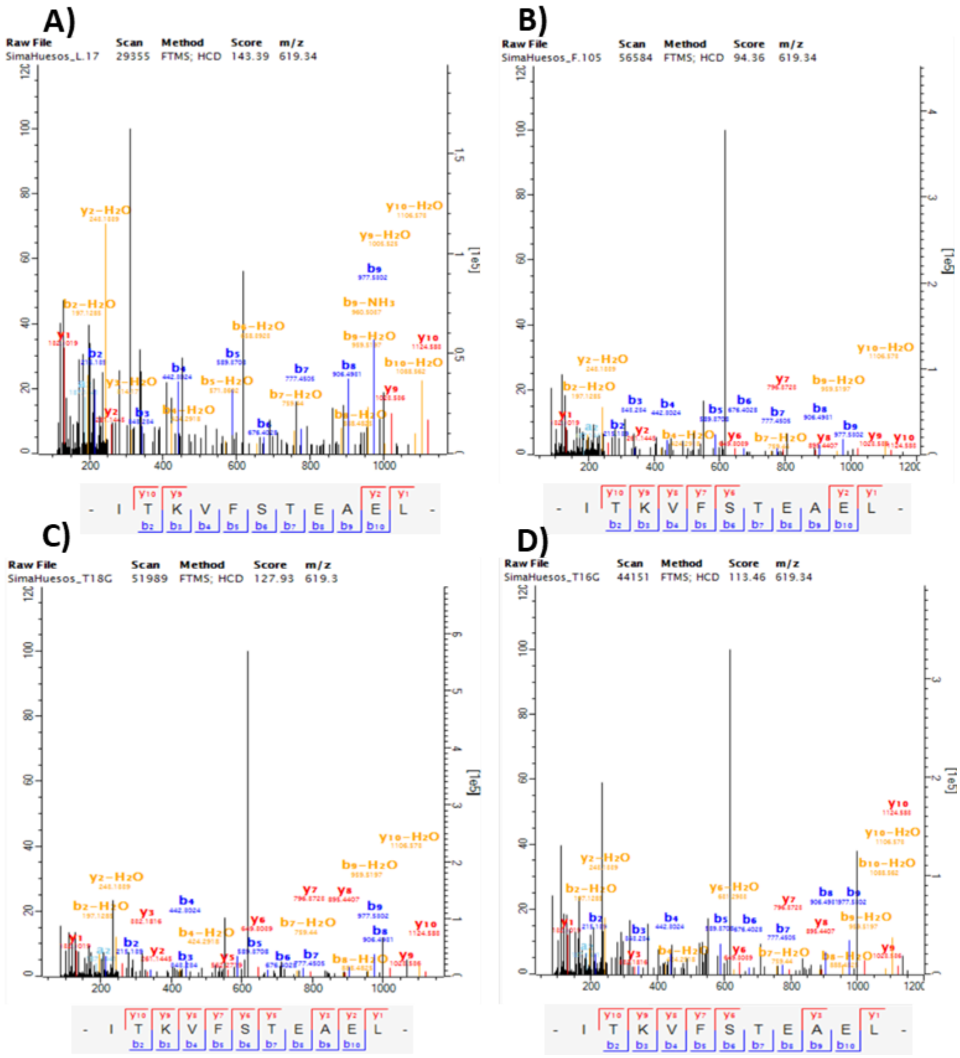


**Figure S20.** Selection of mass spectra and corresponding fragment ions covering SERPINA1-341 obtained with MaxQuant from *Ursus deningeri* specimens. (A) Sample SH.T18.L17. (B) Sample SH.Q20.F105. (C) Sample SH.T18.L75. (D) Sample SH.T16G.

The Monte Cucco *Ursus* samples reveal unusual signatures in SERPINA1. At position 341, both MC.879 and MC.A3 display threonine (T) instead of the conserved asparagine (N) observed in all reference sequences. This variant is supported by overlapping peptides identified with MaxQuant and PEAKS (Fig. S22). In addition, MC.804 exhibits allelic diversity at position 355, with peptides containing lysine (K, ~75%) and methionine (M, ~25%). While K only matches *U. americanus. T*heir co-occurrence suggests heterozygosity or unrecognized allelic variation at this site (Fig. S23). These findings underscore that each Monte Cucco specimen carries a distinct proteomic signature, adding complexity to the evolutionary interpretation of the locus. A further possibility is misattribution of specimens, since *U. spelaeus* is also reported from Monte Cucco. However, the morphometric data do not support a reassignment to this taxon [39].

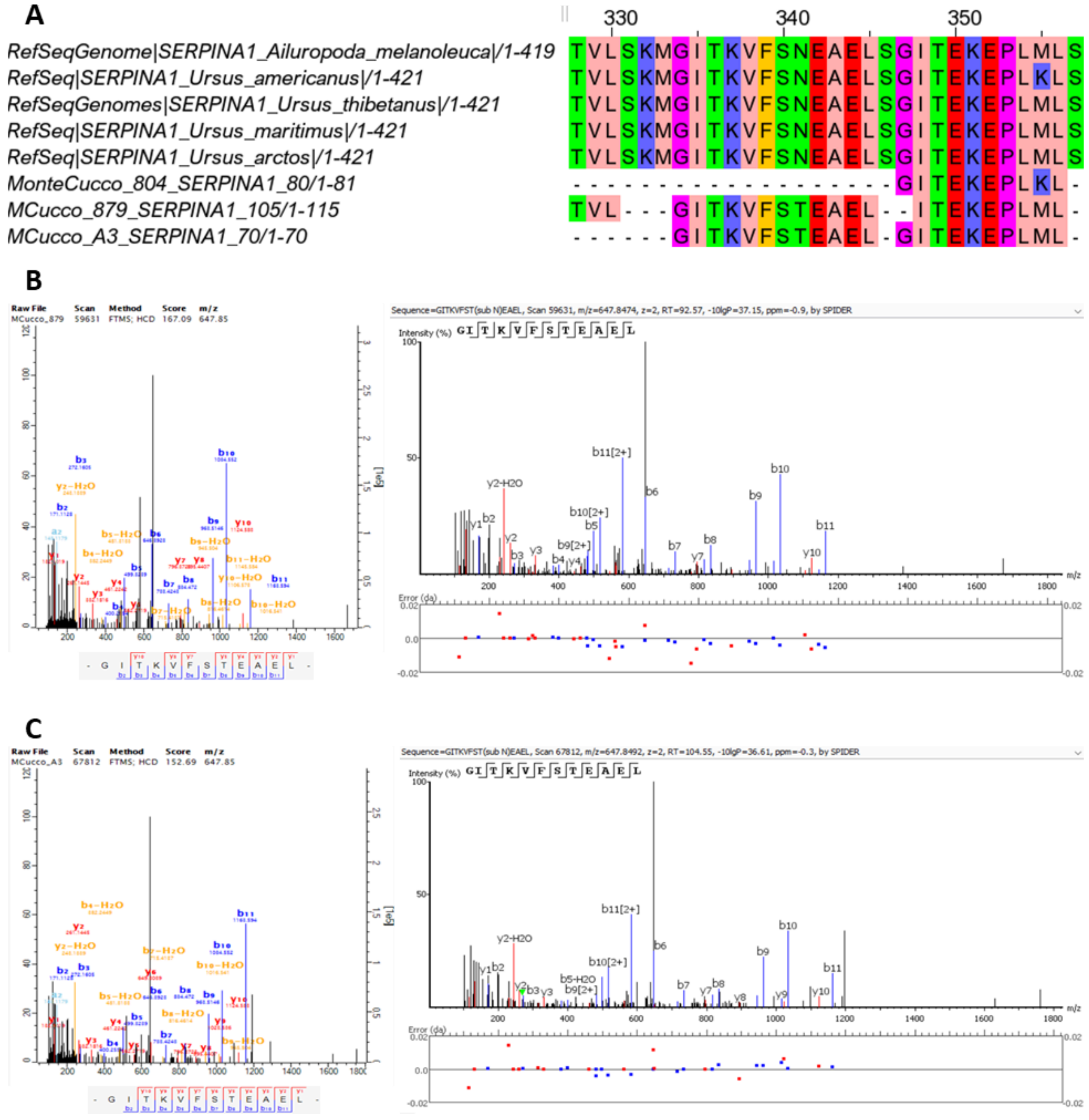


**Figure S21. Variability in SERPINA1 from Monte Cucco specimens.** (A) Multiple sequence alignment of SERPINA1 showing two variable sites (positions 341 and 355) in the Monte Cucco samples compared with the reference dataset. (B) Representative MS/MS spectra from MC.879 *Ursus arctos*, supporting the novel T substitution at position 341, identified with MaxQuant and PEAKS. (C) Equivalent spectra from MC.A3 *Ursus spelaeus* confirming the same substitution, identified with MaxQuant and PEAKS.

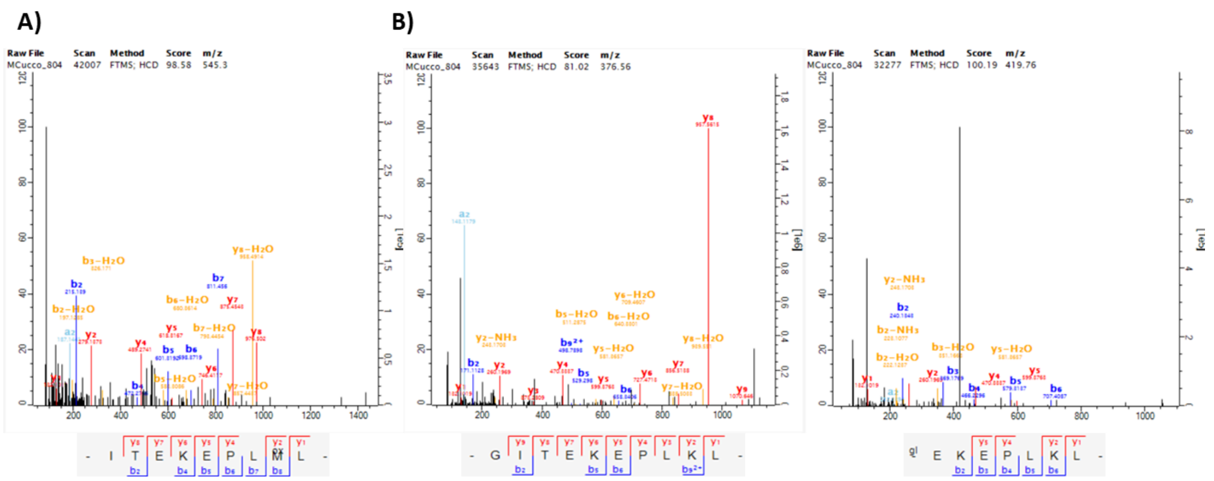


**Figure S22. SERPINA1-355 variation in MC.804.** Representative MS/MS spectra obtained with MaxQuant showing two coexisting variants at position 355: (A) methionine (M) and (B) lysine (K).

##### II.2.4. Validation of SAP ENAM-269

A peptide variant at position ENAM-269 (asparagine (N)→ isoleucine (I)) was consistently detected in seven specimens of *U. deningeri* and *U. spelaeus*, as well as in *U. maritimus*, but not in *U. dolinensis*, where no peptide coverage was recovered at this site. The substitution is supported by multiple spectra independently identified with both MaxQuant and PEAKS, with additional evidence of Q and N deamidation reinforcing the authenticity of the ancient peptides [40].

At the nucleotide level, this change could result from two possible single-base substitutions (AAU→AUU or AAC→AUC), both requiring an A→U transition at the second codon position. The presence of the same substitution in both cave bears and *U. maritimus* may reflect independent parallel evolution driven by similar selective pressures, or alternatively, complex demographic dynamics, given the documented hybridization of *U. arctos* with both *U. maritimus* and *U. spelaeus* [41-43]. The absence of peptide coverage in Early Pleistocene specimens and the potential role of introgression highlight the need for additional molecular data to resolve the evolutionary history of this variant.

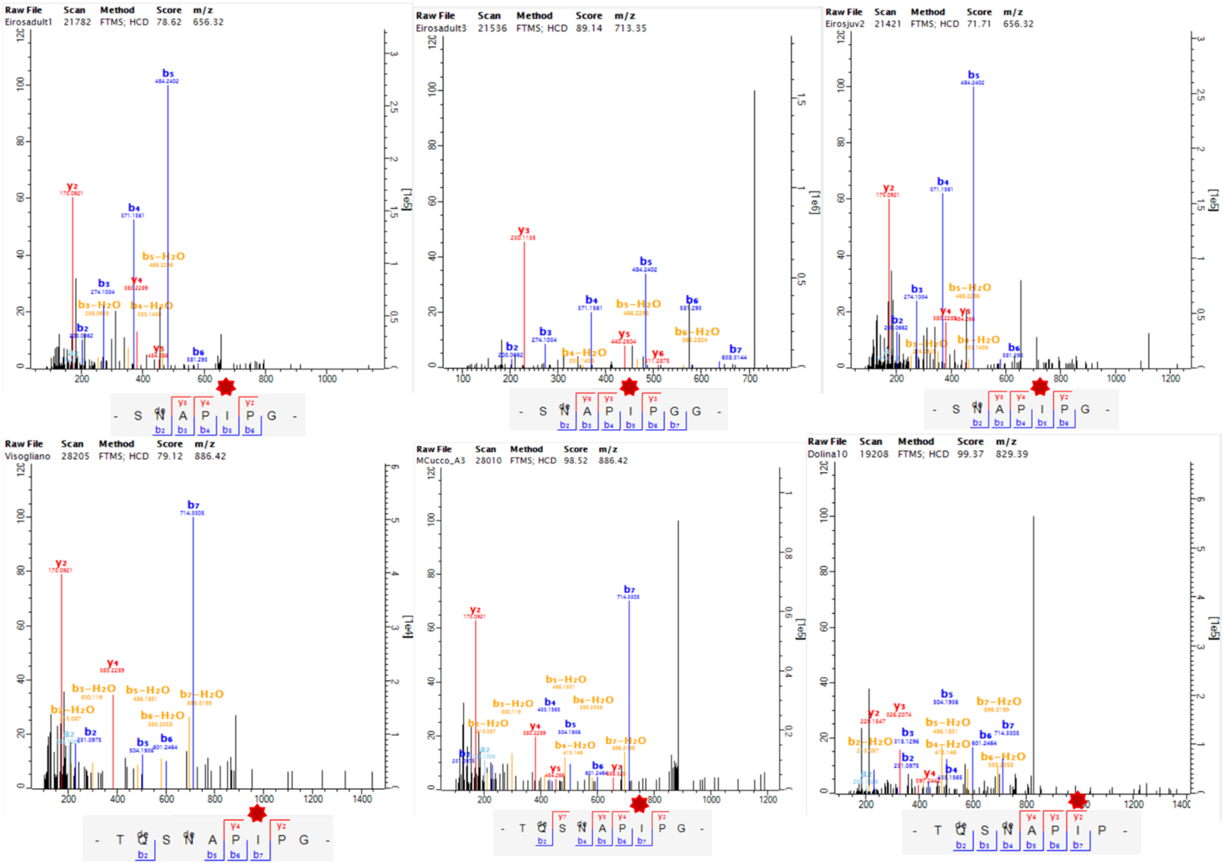

**Figure S23.** Mass spectra and corresponding fragment ions covering ENAM-1328 in several samples from the Middle and Late Pleistocene using MaxQuant. ENAM-1328 position is highlighted by a red asterisk.

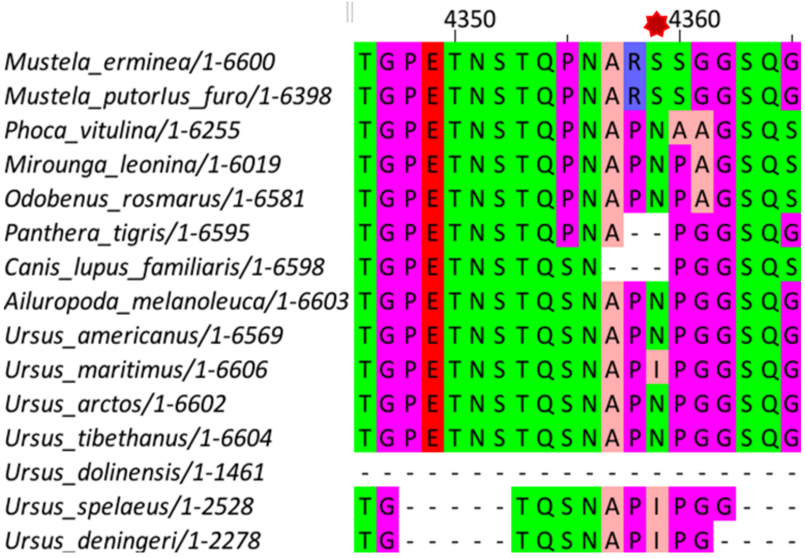


**Figure S24.** Multiple sequence alignment of ENAM across different Carnivora species. ENAM-1328 position is highlighted by a red asterisk.

#### II.3. Expanded Phylogenetic Trees and Node Support

To complement the main phylogenetic reconstructions, we generated additional trees including all available specimens at the individual level.

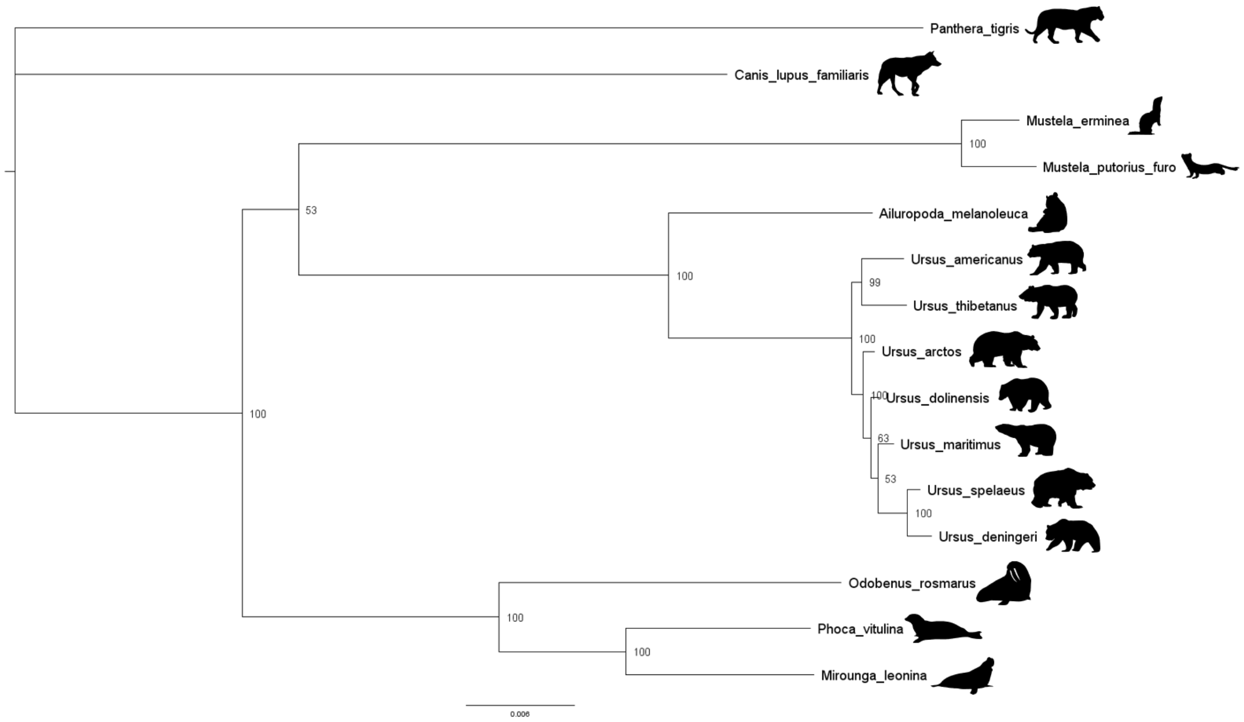


**Figure S25.** Bayesian phylogenetic tree generated using MrBayes and the “Carnivora Dataset”.

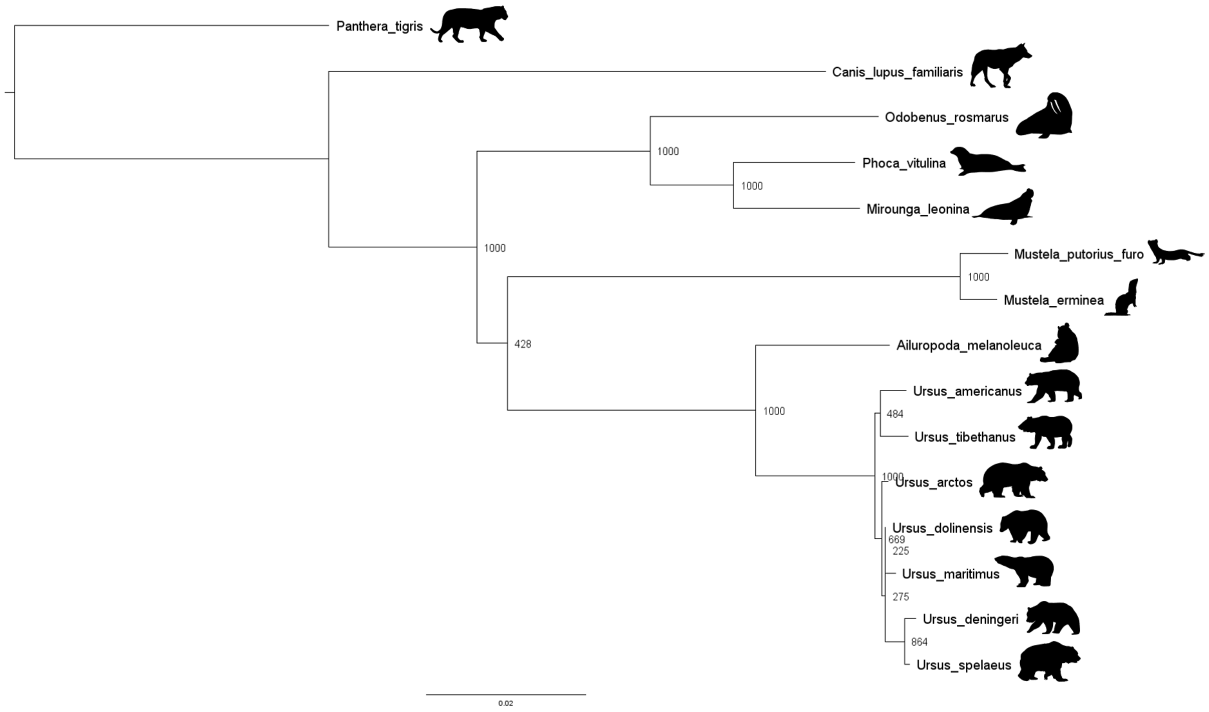


**Figure S26.** Maximum likelihood phylogenetic tree generated using PhyML and the “Carnivora Dataset”.

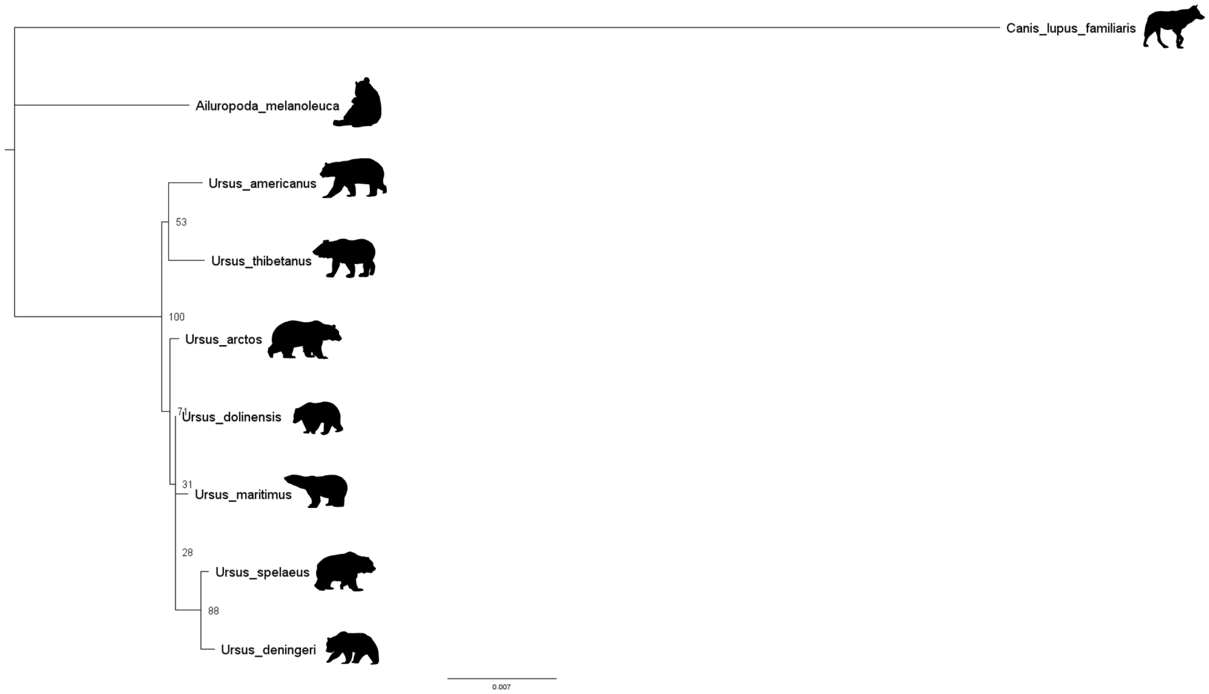


**Figure S27.** Maximum likelihood phylogenetic tree generated using IQTree and the "Ursidae Dataset".

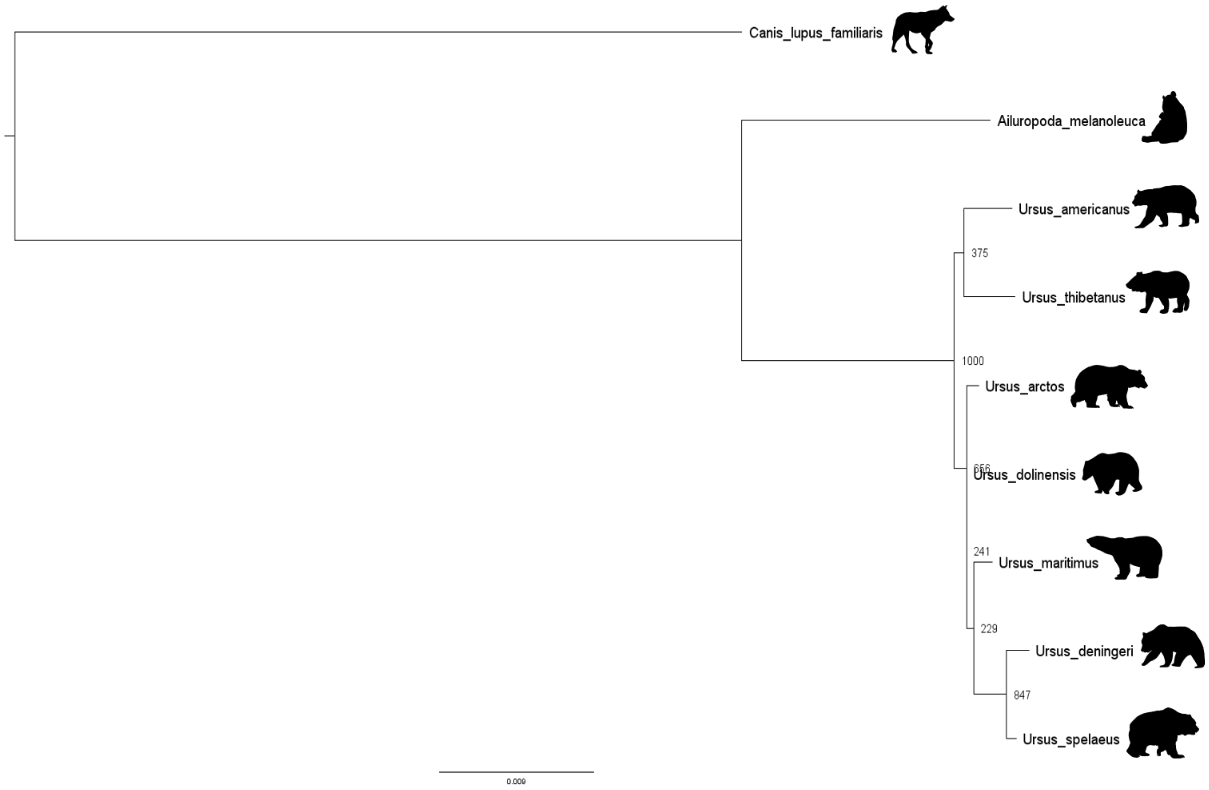


**Figure S28.** Maximum likelihood phylogenetic tree generated using PhyML and the "Ursidae Dataset".

To explore intraspecific variability and clarify ambiguous samples, we generated three additional phylogenetic trees including individual samples rather than grouped taxa. These datasets incorporated specimens assigned to *U. dolinensis* (TD4 from Gran Dolina), *U. deningeri* (Sima de los Huesos, Sala Oseras, Aroeira, Visogliano, Baio), and *U. spelaeus* (Coro Tracito, Cova Eirós, Monte Cucco), as well as *U. arctos* from the Late Pleistocene. Additional material came from Gran Dolina (TD8 and TD10) and Fantasma. Samples from Pirro Nord and Estació Vallparadís were excluded due to extensive fragmentation and limited peptide recovery. These specimens yielded only a few SAPs, insufficient for reliable comparisons. Of the seven most informative positions identified across the dataset, only AMBN-278 was consistently detected in Vallparadís and Pirro Nord, while SERPINA1-355 appeared in a single specimen (EVT12.2). Given the large amount of missing data, including them would have compromised phylogenetic resolution.

Maximum likelihood analyses (IQTree, PhyML) recovered the expected split between arctoid and speloid lineages, consistent with previous studies. Support values varied (IQTree bootstrap = 58/100; PhyML = 224/1000), but both approaches confirmed the separation. As in earlier analyses, *Ursus maritimus* clustered within the speloid lineage, close to the divergence with *U. arctos*, albeit with weak support (IQTree = 34/100; PhyML = 32/1000). Notably, specimen SF30.A.G30 (Sala Fantasma, *Ursus* sp.) grouped with *U. arctos* in both analyses, with moderate support (IQTree = 49/100; PhyML = 104/1000), suggesting shared ancestry.

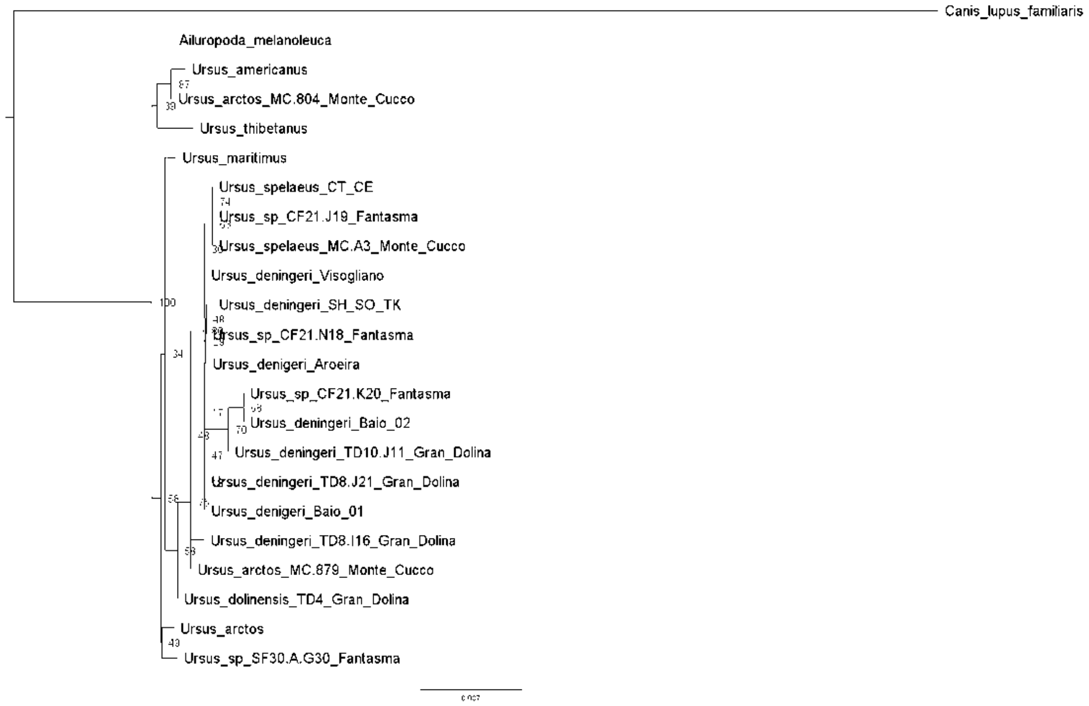


**Figure S29.** Maximum likelihood phylogenetic tree generated using IQTree and the "Ursidae Dataset" including all recovered specimens.

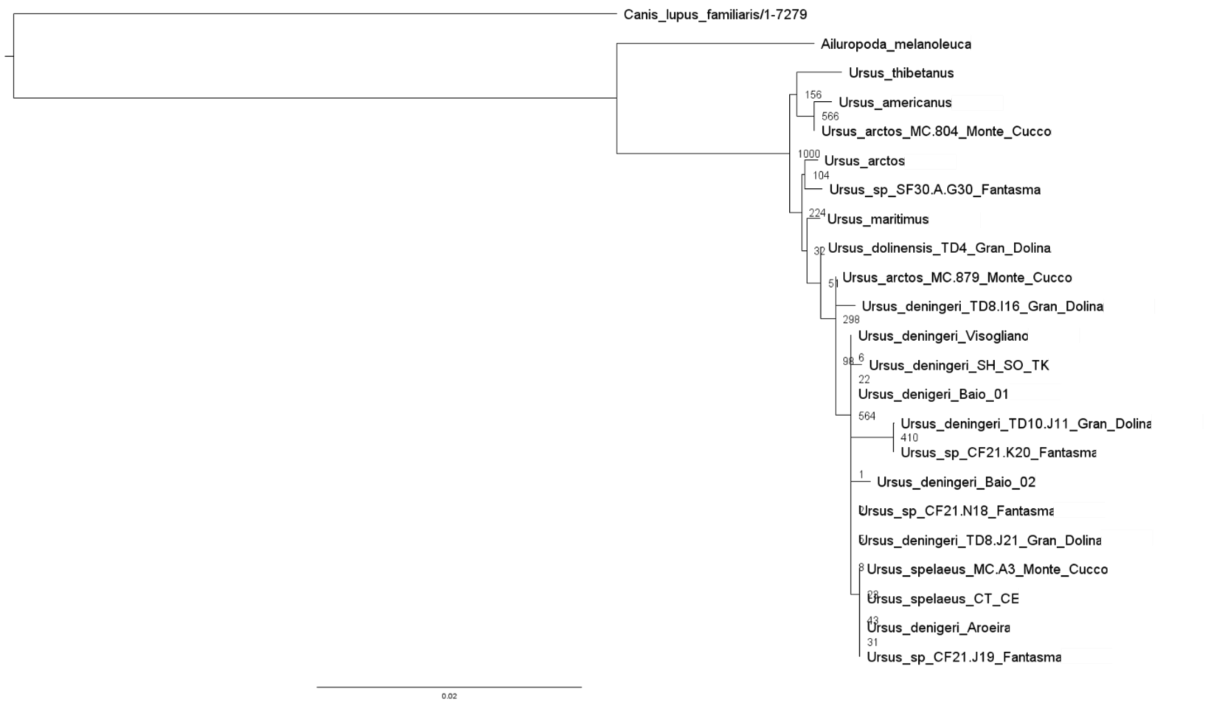


**Figure S30.** Bayesian phylogenetic tree generated using MrBayes and the "Ursidae Dataset" including all recovered specimens.

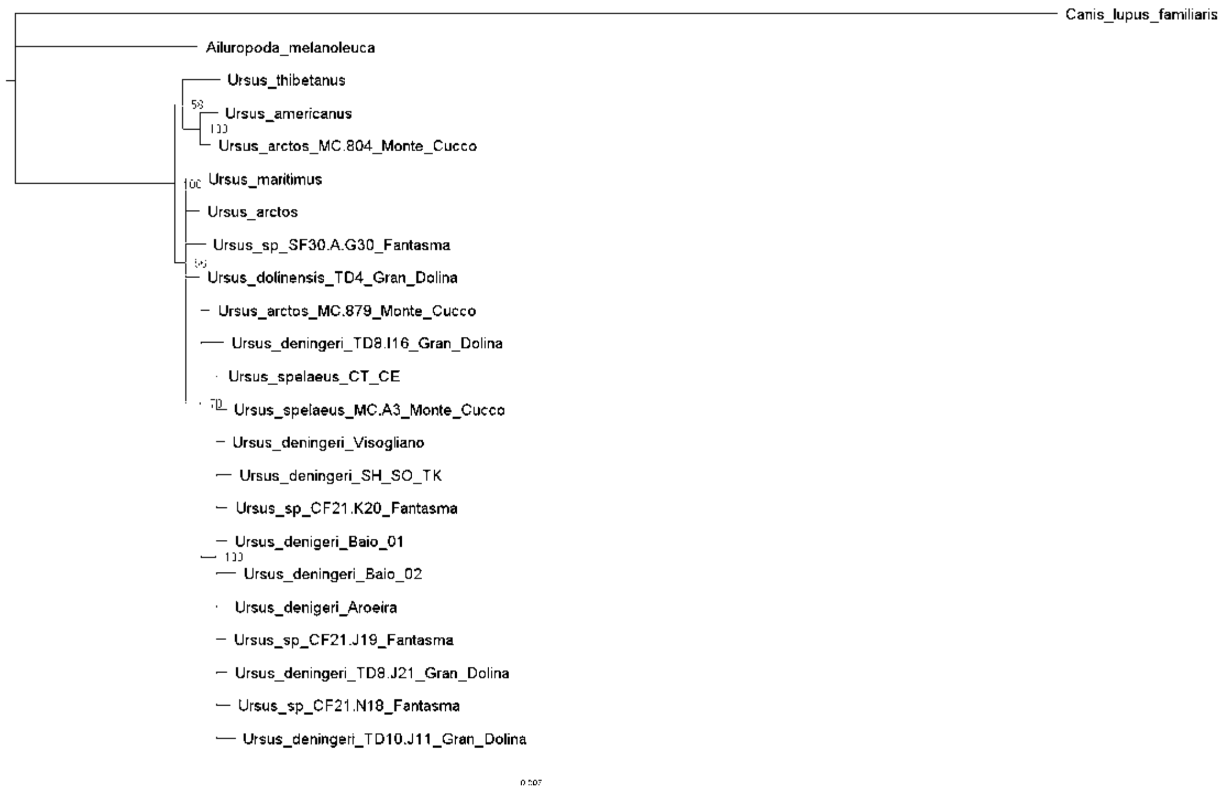


**Figure S31.** Bayesian phylogenetic tree generated using MrBayes and the "Ursidae Dataset" including all recovered specimens.

#### II.4. Sex Determination: Peptides from AMELX/AMELY

The reconstructed *U. arctos* AMELX sequence is 201 amino acids in length, while AMELY spans 182 amino acids, both including an N-terminal signal peptide of 16 residues. Overall sequence identity between isoforms is 89.05%, consistent with previously reported divergence between X- and Y-linked amelogenins in mammals. Despite the high level of conservation, several diagnostic differences distinguish the two isoforms.

At the N-terminal region, AMELX contains an alanine at position 16, whereas AMELY displays a threonine, marking the earliest divergence point between the sequences. A single-residue deletion is observed at position 45, where AMELY lacks a methionine present in AMELX. Further substitutions occur in the central portion of the sequence: at position 101, methionine in AMELX is replaced by valine in AMELY, while at position 104, leucine is replaced by valine. The most striking difference is located in the C-terminal domain, where AMELY exhibits an 18–amino acid deletion spanning positions 138–156, a feature that dramatically reduces the length and potentially alters the biochemical properties of the isoform.

| Position | AMELX residue | AMELY residue | Note |
|---|---|---|---|
| 16 | Ala (A) | Thr (T) | Substitution |
| 45 | Met (M) | – (deletion) | Single-residue deletion |
| 101 | Met (M) | Val (V) | Substitution |
| 104 | Leu (L) | Val (V) | Substitution |
| 138–156 | Present | – (deletion) | 18-aa block deletion |

**Table S2.** Amino acid differences between reconstructed *U. arctos* AMELX and AMELY sequences.

These modifications, particularly the single-residue deletion at position 45 and the substitutions at positions 101 and 104, generate robust isoform-specific peptide signatures that are repeatedly detected in our dataset. Multiple high-quality spectra were recovered for these regions in both AMELX and the Y-linked isoform, providing strong support for the reliability of the assignments.

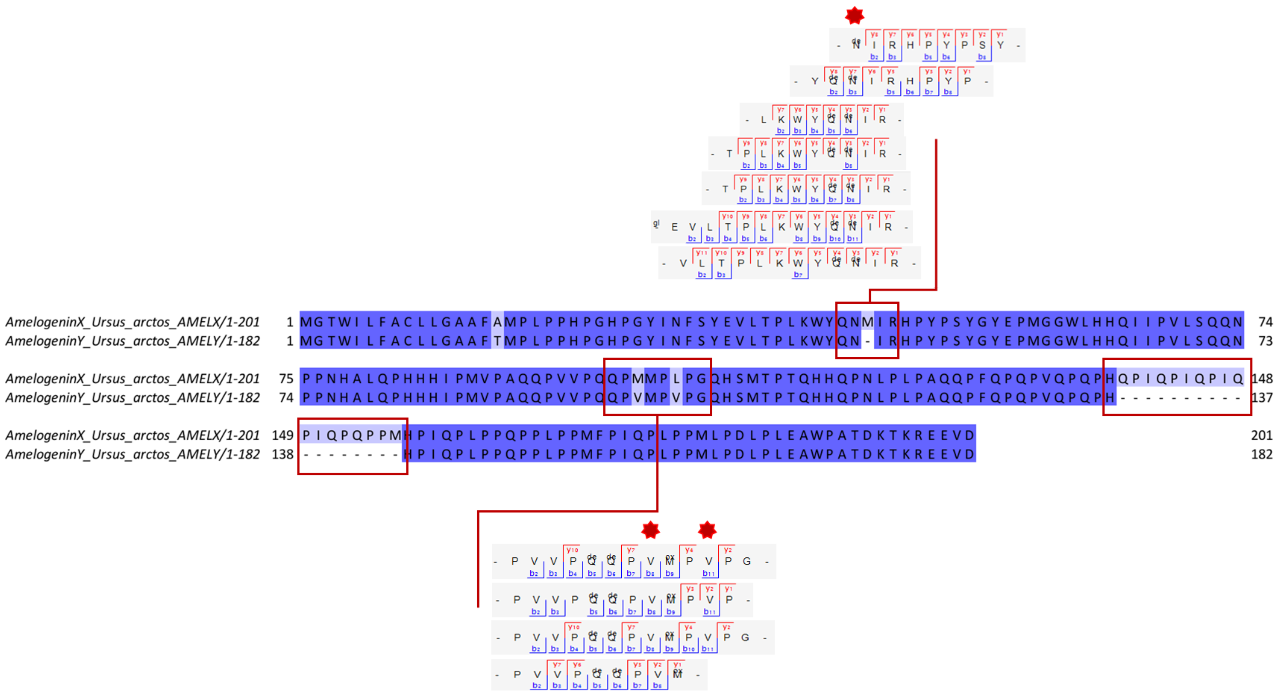


**Figure S32. Alignment of reconstructed *Ursus arctos* AMELX and AMELY sequences.** Amino acid alignment highlighting diagnostic differences between X- and Y-linked isoforms. Positions 16, 45, 101, and 104 display single-residue substitutions or deletions, while AMELY exhibits an additional 18–amino acid deletion (positions 138–156). Regions surrounding positions 45, 101, and 104 yielded multiple high-quality spectra for both isoforms, making them particularly informative for sex determination. Diagnostic ion series from representative peptides are shown below, illustrating the MS/MS fragmentation patterns that confirm the presence of the Y-specific isoform in male specimens.

#### SI VI. Supplementary References


1. W. V. Koenigswald, W. A. Clemens, Levels of Complexity in the Microstructure of Mammalian Enamel and Their Application in Studies of Systematics. *Scanning Microscopy* **6** (1992).

2. C. Stefen, Enamel structure of arctoid carnivora: Amphicyonidae, Ursidae, Procyonidae, and Mustelidae. *Journal of Mammalogy* **82**, 450–462 (2001).

3. T. Wiszniowska, *et al.*, Dental enamel structure in fossil bears *Ursus spelaeus* and *U. wenzensis* (= *minimus*) in comparison to selected representatives of other Carnivora. *Morphology and systematics of fossil vertebrates. DN Publisher, Wrocław* 125 (2010).

4. P. Mackiewicz, *et al.*, "Analysis of dental enamel thickness in bears with special attention to *Ursus spelaeus* and *U. wenzensis* (= *minimus*) in comparison to selected representatives of mammals" in (DN Publisher, 2010).

5. J. Rappsilber, M. Mann, Y. Ishihama, Protocol for micro-purification, enrichment, pre-fractionation and storage of peptides for proteomics using StageTips. *Nature protocols* **2**, 1896–1906 (2007).

6. C. Chiva, *et al.*, QCloud: A cloud-based quality control system for mass spectrometry-based proteomics laboratories. *PloS one* **13**, e0189209 (2018).

7. I. Patramanis, J. Ramos-Madrigal, E. Cappellini, F. Racimo, PaleoProPhyler: a reproducible pipeline for phylogenetic inference using ancient proteins. *Peer Community Journal* **3** (2023).

8. T. Kishida, M. Ohashi, Y. Komatsu, Genetic diversity and population history of the Japanese black bear ( *Ursus thibetanus japonicus* ) based on the genome-wide analyses. *Ecological Research* **37**, 647–657 (2022).

9. S. Chen, Y. Zhou, Y. Chen, J. Gu, fastp: an ultra-fast all-in-one FASTQ preprocessor. *Bioinformatics* **34**, i884–i890 (2018).

10. E. E. Armstrong, *et al.*, A Beary Good Genome: Haplotype-Resolved, Chromosome-Level Assembly of the Brown Bear (*Ursus arctos*). *Genome Biology and Evolution* **14**, evac125 (2022).

11. H. Li, Aligning sequence reads, clone sequences and assembly contigs with BWA-MEM. *arXiv preprint arXiv:1303.3997* (2013).

12. G. Tischler, S. Leonard, biobambam: tools for read pair collation based algorithms on BAM files. *Source Code Biol Med* **9**, 13 (2014).

13. A. McKenna, *et al.*, The Genome Analysis Toolkit: A MapReduce framework for analyzing next-generation DNA sequencing data. *Genome Res.* **20**, 1297–1303 (2010).

14. R. Fong-Zazueta, *et al.*, Phylogenetic Signal in Primate Tooth Enamel Proteins and its Relevance for Paleoproteomics. *Genome Biol Evol* **17**, evaf007 (2025).

15. A. Barlow, *et al.*, Partial genomic survival of cave bears in living brown bears. *Nat Ecol Evol* **2**, 1563–1570 (2018).

16. G. G. Fortes, *et al.*, Ancient DNA reveals differences in behaviour and sociality between brown bears and extinct cave bears. *Molecular Ecology* **25**, 4907–4918 (2016).

17. H. Li, R. Durbin, Fast and accurate short read alignment with Burrows-Wheeler transform. *Bioinformatics* **25**, 1754–1760 (2009).

18. T. S. Korneliussen, A. Albrechtsen, R. Nielsen, ANGSD: Analysis of Next Generation Sequencing Data. *BMC Bioinformatics* **15**, 356 (2014).

19. F. Madeira, *et al.*, Search and sequence analysis tools services from EMBL-EBI in 2022. *Nucleic acids research* **50**, W276–W279 (2022).

20. E. Gasteiger, *et al.*, ExPASy: the proteomics server for in-depth protein knowledge and analysis. *Nucleic acids research* **31**, 3784–3788 (2003).

21. E. C. Salido, P. H. Yen, K. Koprivnikar, L.-C. Yu, L. J. Shapiro, The human enamel protein gene amelogenin is expressed from both the X and the Y chromosomes. *American journal of human genetics* **50**, 303 (1992).

22. J. Cox, M. Mann, MaxQuant enables high peptide identification rates, individualized ppb-range mass accuracies and proteome-wide protein quantification. *Nature biotechnology* **26**, 1367–1372 (2008).

23. J. Cox, Prediction of peptide mass spectral libraries with machine learning. *Nature Biotechnology* **41**, 33–43 (2023).

24. J. Zhang, *et al.*, PEAKS DB: de novo sequencing assisted database search for sensitive and accurate peptide identification. *Molecular & cellular proteomics* **11** (2012).

.25. Y. Han, B. Ma, K. Zhang, SPIDER: software for protein identification from sequence tags with de novo sequencing error in *Proceedings. 2004 IEEE Computational Systems Bioinformatics Conference, 2004. CSB 2004.*, (2004), pp. 206–215.

26. P. Gutenbrunner, P. Kyriakidou, F. Welker, J. Cox, Spectrum graph-based de-novo sequencing algorithm MaxNovo achieves high peptide identification rates in collisional dissociation MS/MS spectra. *bioRxiv* 2021–09 (2021).

27. P. P. Madupe, *et al.*, Enamel proteins reveal biological sex and genetic variability in southern African *Paranthropus*. *Science* **388**, 969–973 (2025).

28. F. Welker, Elucidation of cross-species proteomic effects in human and hominin bone proteome identification through a bioinformatics experiment. *BMC Evol Biol* **18**, 23 (2018).

29. A. J. Taurozzi, *et al.*, Deep-time phylogenetic inference by paleoproteomic analysis of dental enamel. *Nat Protoc* **19**, 2085–2116 (2024).

30. E. Cappellini, et al., Early Pleistocene enamel proteome from Dmanisi resolves *Stephanorhinus* phylogeny. Nature 574, 103–107 (2019).

31. J. Hendy, Ancient protein analysis in archaeology. *Sci. Adv.* **7**, eabb9314 (2021).

32. F. Welker, *et al.*, The dental proteome of *Homo antecessor*. *Nature* **580**, 235–238 (2020).

33. K. Katoh, K. Misawa, K. Kuma, T. Miyata, MAFFT: a novel method for rapid multiple sequence alignment based on fast Fourier transform. *Nucleic Acids Research* **30**, 3059–3066 (2002).

34. J. Castresana, Selection of Conserved Blocks from Multiple Alignments for Their Use in Phylogenetic Analysis. *Molecular Biology and Evolution* **17**, 540–552 (2000).

35. L.-T. Nguyen, H. A. Schmidt, A. Von Haeseler, B. Q. Minh, IQ-TREE: a fast and effective stochastic algorithm for estimating maximum-likelihood phylogenies. *Molecular biology and evolution* **32**, 268–274 (2015).

36. S. Kalyaanamoorthy, B. Q. Minh, T. K. Wong, A. Von Haeseler, L. S. Jermiin, ModelFinder: fast model selection for accurate phylogenetic estimates. *Nature methods* **14**, 587–589 (2017).

37. S. Guindon, *et al.*, New algorithms and methods to estimate maximum-likelihood phylogenies: assessing the performance of PhyML 3.0. *Systematic biology* **59**, 307–321 (2010).

38. F. Ronquist, *et al.*, MrBayes 3.2: Efficient Bayesian Phylogenetic Inference and Model Choice Across a Large Model Space. *Systematic Biology* **61**, 539–542 (2012).

39. L. Capasso Barbato, M. R. Minieri, C. Petronio, A. Vigna Taglianti, Strutture dentarie di *Ursus arctos* e di *Ursus spelaeu*s della Grotta di Monte Cucco (Sigillo, Perugia, Italia). *Bollettino della Società paleontologica italiana* **29** (1990).

40. E. R. Schroeter, T. P. Cleland, Glutamine deamidation: an indicator of antiquity, or preservational quality? *Rapid Communications in Mass Spectrometry* **30**, 251–255 (2016).

41. J. D. Pongracz, D. Paetkau, M. Branigan, E. Richardson, Recent Hybridization between a Polar Bear and Grizzly Bears in the Canadian Arctic. *Arctic* **70**, 151–160 (2017).

42. J. A. Cahill, *et al.*, Genomic evidence of geographically widespread effect of gene flow from polar bears into brown bears. *Molecular Ecology* **24**, 1205–1217 (2015).

43. J. A. Cahill, *et al.*, Genomic evidence of widespread admixture from polar bears into brown bears during the last ice age. *Molecular Biology and Evolution* **35**, 1120–1129 (2018).
